# Adult *Hox* gene expression promotes periosteal stem cell maintenance and mediates reprogramming in a regionally restricted manner

**DOI:** 10.1101/2022.05.15.492027

**Authors:** Kevin Leclerc, Lindsey H. Remark, Malissa Ramsukh, Anne Marie Josephson, Sophie M. Morgani, Laura Palma, Paulo EL Parente, Sooyeon Lee, Emma Muiños Lopez, Philipp Leucht

## Abstract

Periosteal stem and progenitor cells are pivotal to the growth and lifelong turnover of bone and underpin its capacity to regenerate. Adjusting the potency of this cell population will therefore be critical to the successful generation and application of new bone repair therapies. Following their role in patterning the embryonic skeleton, *Hox* genes remain regionally expressed in mesenchymal stromal cell populations of the adult skeleton. Here we show that *Hoxa10* is most expressed in the most uncommitted periosteal stem cell and that *Hox* maintains these skeletal stem cells in a multipotential, uncommitted state, thereby preventing their differentiation into bone. We demonstrate that *Hoxa10* mediates the reprogramming of periosteal progenitors towards a stem cell state with greater self-renewal capacity and also establish that region-specific *Hox* genes mediate cell reprogramming in distinct anatomical regions, demonstrating the continued functional relevance of the embryonic *Hox* profile in adult stem cells. Together, our data describe a master regulator role of *Hox* in skeletal stem and progenitor cells and help provide insight into the development of cell-based therapies for treatment of at-risk bone fractures and other bone-related ailments.

## Introduction

Bone homeostasis and repair is mediated by skeletal stem cells (SSCs) that self-renew and differentiate into the major skeletal/mesenchymal lineages (stromal, osteo-, chondro-, and adipo-lineages) (Zhou, Yue et al. 2014, Yue, Zhou et al. 2016, Ambrosi, Longaker et al. 2019). SSCs harbor massive therapeutic potential as an unlimited source of differentiated cells to replace those lost and damaged in injury and disease. However, during aging, they are relatively rare and the molecular and genetic mechanisms that regulate stem cell number and function are still largely unknown.

Skeletal stem and progenitor cells reside in the bone marrow of long bones, and also on the inner and outer surfaces of cortical bone. It was recently shown that the periosteum, a layer of mesenchymal cells that lines the outer cortex of the bone, is highly enriched for SSCs that are transcriptionally distinct from stem cells from other regions (Duchamp de Lageneste, Julien et al. 2018). Although periosteal cells have higher regenerative capacity than mesenchymal cells derived from bone marrow, they remain poorly characterized. Recent evidence suggests that periosteal stem and progenitor cells (PSPCs) are the main contributor to the fracture callus after long bone fractures (Colnot, Zhang et al. 2012, Ferretti and Mattioli-Belmonte 2014, Roberts, van Gastel et al. 2015, Duchamp de Lageneste, Julien et al. 2018), pinpointing this population as an important therapeutic target to improve fracture healing in patients that are prone to developing a nonunion – such as aged patients (Nilsson and Edwards 1969, Nieminen, Nurmi et al. 1981, Gruber, Koch et al. 2006, Kwong and Harris 2008) or patients with high-energy injuries and accompanying soft tissue loss (Harris, Althausen et al. 2009, Cheng, Vantucci et al. 2021).

Homeobox (Hox) genes are evolutionarily conserved transcription factors that are master regulators of positional identity and cell fate specification during embryonic development (Deschamps and van Nes 2005). The mouse and human genomes contain 39 *Hox* genes, which are grouped into four clusters, *Hoxa*, *Hoxb*, *Hoxc*, and *Hoxd*, positioned on four separate chromosomes in 13 paralogs (Izpisua-Belmonte, Falkenstein et al. 1991, Krumlauf 1994). During development, overlapping patterns of *Hox* gene activity assign a segmental identity to each anatomic body part, which culminates in the creation of a complex tissue, organ or organism. While this genetic blueprint is essential to establish the body plan of an organism, its function in adulthood is unclear. *Hox* genes are expressed during homeostasis and tissue repair (Pineault, Helgason et al. 2002, Leucht, Kim et al. 2008, Gerber, Murawala et al. 2018, Bradaschia-Correa, Leclerc et al. 2019, Lin, Gerber et al. 2021), suggesting that the Hox code may also be required for successful regeneration and could similarly impart positional information and, in skeletal tissues, *Hoxa11*-expressing cells comprise a stem/progenitor cell population necessary for successful fracture healing of the ulna (Rux, Song et al. 2016, Song, Pineault et al. 2020). However, the precise function of *Hox* genes in adulthood is yet unknown and, surprisingly, little is known regarding the regional specificity of adult *Hox* gene function.

Here we identify *Hox* genes as key regulators of skeletal stem cell maintenance. We show that *Hox* deficiency leads to a decrease in stem cell number and proliferation, and an increase in their propensity to undergo spontaneous differentiation toward osteogenic, adipogenic, and chondrogenic fates. Conversely, an increase in *Hoxa10* expression reduces PSPC differentiation potential and increases the self-renewal capacity of the stem cell compartment. During differentiation, SSCs give rise to a larger number of fate-restricted progenitors that have limited lineage potential and lifespan (Chan, Seo et al. 2015, Debnath, Yallowitz et al. 2018). Here, we also uncover the ability of *Hox* genes to drive periosteal progenitor reprogramming to a more uncommitted bona fide stem cell state and further establish that the reprogramming capacity of specific *Hox* genes is restricted to the anatomical regions that reflect their embryonic expression pattern. Harvesting and reprogramming more prevalent progenitor populations may represent an innovative source of SSCs and a promising strategy to combat bone deficiencies, such as the aged-associated bone loss driven by a decline in SSC number and function (Josephson, Bradaschia-Correa et al. 2019).

## Results

### Hox gene expression is enriched in skeletal stem/ progenitor cells and declines with age

*Hox* genes regulate morphogenesis and regeneration in multiple organisms (Gardiner and Bryant 1996, Orii, Kato et al. 1999, Nogi and Watanabe 2001, Christen, Beck et al. 2003, Thummel, Ju et al. 2007). In several vertebrate organs, *Hox* gene expression marks subpopulations of mesenchymal stem cells (Hwang, Seok et al. 2009, Liedtke, Buchheiser et al. 2010, Rux, Song et al. 2016, Bradaschia-Correa, Leclerc et al. 2019) that mediate tissue repair. To investigate the role of *Hox* genes within the adult skeleton, we first analyzed their expression within LEPR^+^ skeletal stem and progenitor cells (SSPCs) and the microenvironment – comprising differentiated skeletal lineages, CD45^+^ hematopoietic, CD31^+^ erythroid, and TER119^+^ endothelial cells – using our previously generated RNA-sequencing dataset of hindlimb skeletal elements (Josephson, Bradaschia-Correa et al. 2019). Previous reports showed that *Hoxa11* marks a population of bone marrow stem cells (Rux, Song et al. 2016). Our analysis revealed that all Hox family members are enriched in SSPCs relative to the microenvironment (**Fig. 1A)**, suggesting that they play a crucial role in stem cell function. Specifically, *Hoxa10* was most highly expressed and enriched 50-fold in skeletal stem cells compared to cells of the microenvironment (*p* = 0.0043). SSPC number and function decline with age (Josephson, Bradaschia-Correa et al. 2019, Ambrosi, Marecic et al. 2021), leading to a reduction in the regenerative capacity of bone. Intriguingly, using our RNA sequencing dataset (Josephson, Bradaschia-Correa et al. 2019), we noted that the majority of *Hox* genes were downregulated in LEPR^+^ SSPCs harvested from the hindlimbs of middle-aged (52-week-old) mice when compared to those of young (12-week-old) (**Fig. 1B**). This result was confirmed in bone marrow samples harvested from the fracture sites of young (19-39 years-old) and aged (61-86 years-old) human patients (including 7 males and 8 females) where the older cohort of patients displayed about half of the *Hoxa10* levels expressed by the younger cohort by qRT-PCR (**Fig. 1C**). We further corroborated these data in the periosteal compartment of aged mice, which show a large reduction in the frequency of CD49f^low^CD51^low^:CD200^+^CD105^−^ periosteal stem cells as defined by *Debnath et al.* (Debnath, Yallowitz et al. 2018) (**Fig. 1D**) and display significantly less *Hoxa10* expression by qRT-PCR when compared to young mice (**Fig. 1E**). Altogether these data indicate that *Hox* expression is associated with functional skeletal stem cells.

**Figure 1.**
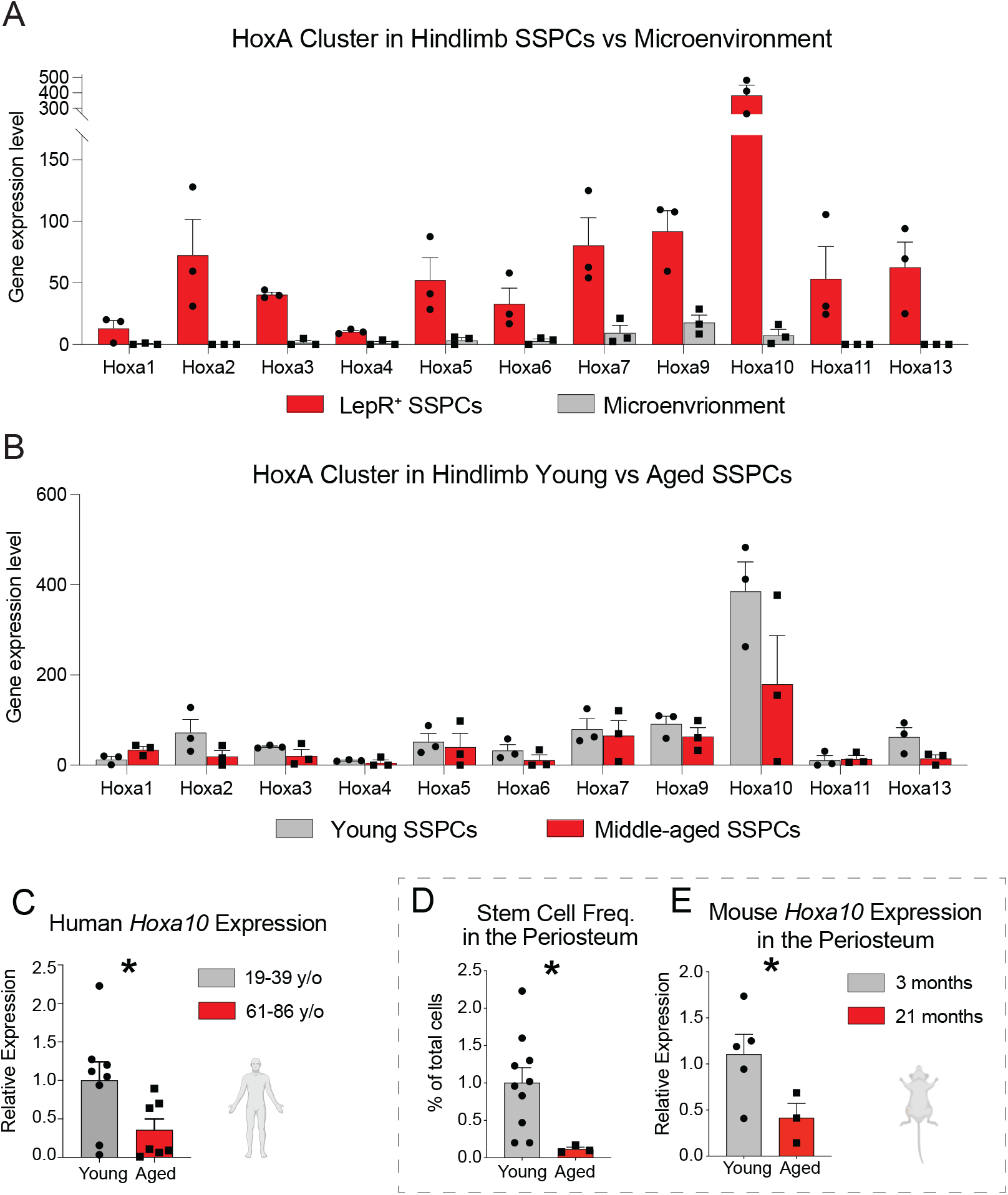
Hox gene expression is enriched in skeletal stem/ progenitor cells and declines with age. (**A**) RNA-sequencing revealed the gene expression pattern for 11 HoxA cluster genes in CD45^−^Ter119^−^CD31^−^LepR+ SSPCs or cells of the microenvironment harvested from 12-week-old, freshly-isolated tibiae and femurs. HoxA genes are highly enriched in the SSPC population and *Hoxa10*, with the most normalized reads, is the most highly expressed. *n* = 3. (**B**) RNA-sequencing determined the gene expression levels of HoxA cluster genes in young, 12-week-old CD45^−^Ter119^−^ CD31^−^LepR+ SSPCs versus those of middle-aged, 52-week-old SSPCs. *n* = 3. (**C**) The relative expression of Hoxa10 in bone marrow samples harvested from the fracture sites of young (18-39 years-old) and aged (61-86 years-old) human patients, as measured by qRT-PCR. *n* = 8 (young), *n* = 7 (aged). (**D** and **E**) When tibial periosteal cell were harvested from young (3-month-old) and aged (21-month-old) mice, flow cytometry revealed the frequency of 6C3^−^CD90^−^CD49f^low^CD51^low^CD200^+^CD105^−^ periosteal stem cells (**D**) and qRT-PCR determined the relative expression of Hoxa10 (**E**). *n* = 10 (young, flow cytometry), *n* = 5 (young, qRT-PCR), *n* = 3 (aged). \**p* < 0.05. Two tailed Student’s t-test. Error bars are SEM.

### Hoxa10 is the most highly expressed Hox gene in tibial periosteal cells

*Hox* gene clusters display spatial collinearity, an increase in the expression of sequential *Hox* genes as development proceeds along the anterior-posterior (A-P) axis. Several *Hox* genes are co-activated at a given position along the A-P axis with the most highly expressed representing the key regulator within that region (Izpisua-Belmonte, Falkenstein et al. 1991, Papageorgiou 2012, Darbellay, Bochaton et al. 2019). We, therefore, hypothesized that *Hox* expression in adult tissues correlates with essential functions. To quantify *Hox* gene expression in skeletal stem cells and progenitors during homeostasis and assess whether this recapitulates the embryonic pattern, we performed bulk RNA sequencing and Nanostring nCounter^®^ gene expression technology on cells isolated from adult tibiae. While numerous skeletal stem cell populations have been described (Costa, Eiro et al. 2021, Mabuchi, Okawara et al. 2021), we chose to focus here on periosteal stem cells as they demonstrate the highest regenerative capacity (Colnot, Zhang et al. 2012, Ferretti and Mattioli-Belmonte 2014, Roberts, van Gastel et al. 2015, Duchamp de Lageneste, Julien et al. 2018). Both analyses indicated that *Hoxa10* is the most highly expressed family member in the adult tibial periosteum (**Fig. 2A, Supplementary Fig. 1A, B**). To conduct a more detailed analysis of *Hoxa10*’s expression pattern within the periosteal compartment, we next separated periosteal cells into periosteal stem cells (PSC) (CD49f^low^CD51^low^:CD200^+^CD105^−^), periosteal progenitor 1 cells (PP1) (CD49f^low^CD51^low^:CD200^−^CD105^−^) and periosteal progenitor 2 cells (PP2) (CD49f^low^CD51^low^:CD105^+^CD200^variable^) using flow cytometry, according to *Debnath et al*. (Debnath, Yallowitz et al. 2018) (**Fig. 2B, C, and Supplementary Fig. 1C**). Using qRT-PCR, we observed high *Hoxa10* expression in PSCs, the most uncommitted compartment, and lower expression in PP1 and PP2 cells (**Fig. 2D**), which are more lineage-restricted (Debnath, Yallowitz et al. 2018). This is consistent with studies in the hematopoietic system, endometrium, and craniofacial skeleton, which show that *Hoxa10* is expressed by adult cells with high plasticity (Kanzler, Kuschert et al. 1998, Creuzet, Couly et al. 2002, Magnusson, Brun et al. 2007, Zanatta, Rocha et al. 2010).

**Figure 2.**
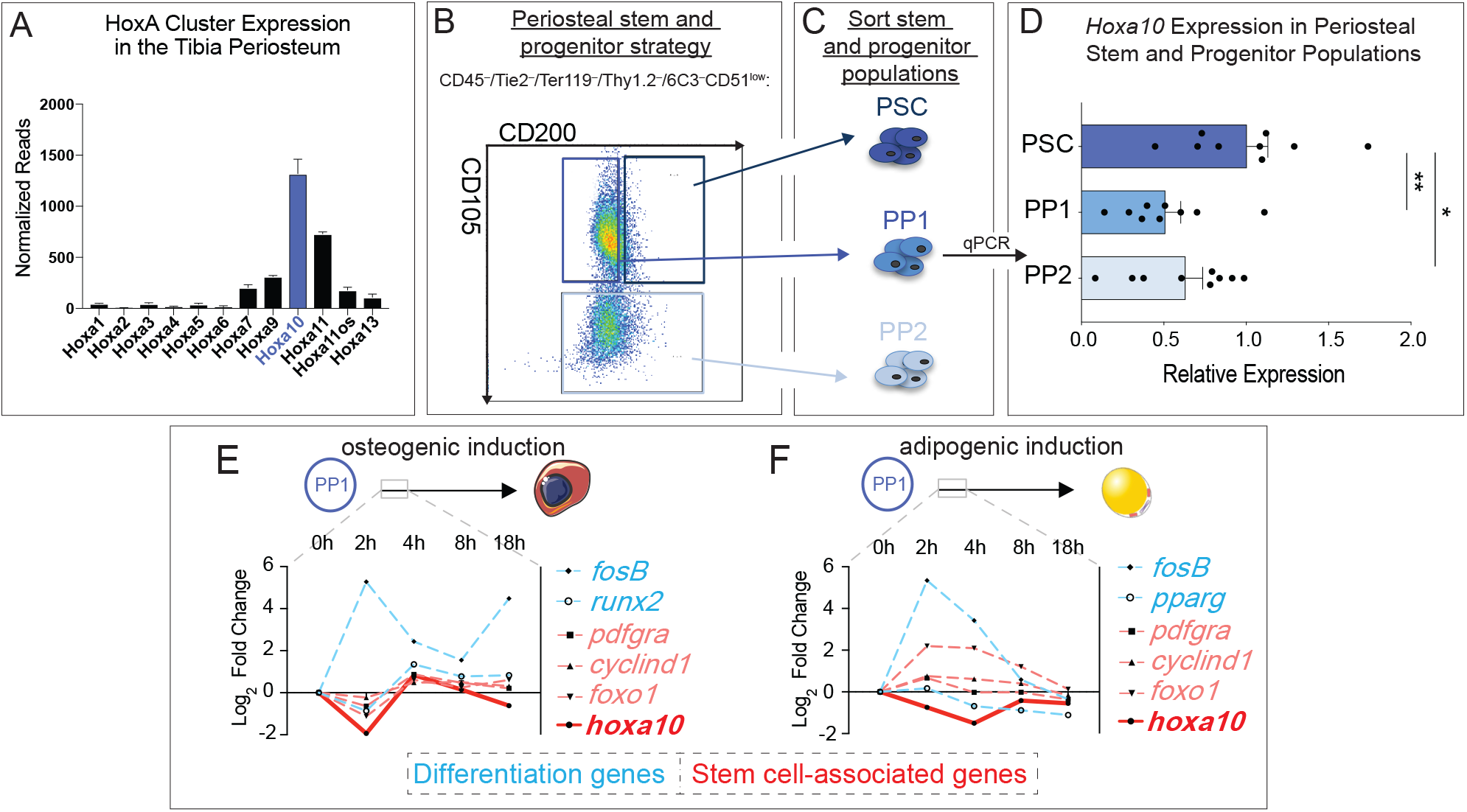
Hoxa10 is the most highly expressed Hox gene in tibial periosteal cells. (**A**) RNAseq gene expression data of the HoxA cluster derived from 12-week-old tibial periosteal cells. (**B**) Sample FACS plot of periosteal stem and progenitor cells as defined by *Debnath et al., 2018* and (**C**) strategy for isolating periosteal stem cells (PSC), periosteal progenitor 1 (PP1) cells, and periosteal progenitor 2 (PP2) cells. (**D**) The relative gene expression of *Hoxa10* in freshly isolated stem and progenitor populations of tibial periosteum as measured by qRT-PCR. *n* = 3 mice. (**E,F**) The gene expression of multiple skeletal stem cell and differentiation genes during an 18h *in vitro* time course of osteogenic (**E**) or adipogenic (**F**) induction relative to growth media controls. *n* = 3. \**p* < 0.05, \*\**p* < 0.01. Two tailed Student’s t-test. Error bars are SEM.

### Hoxa10 is among the first genes to be downregulated at the initiation of differentiation

Stem cell differentiation is often a two-step process that involves first dismantling the gene regulatory network that maintains self-renewal before subsequently upregulating a lineage-specific transcriptional program (Bardin, Perdigoto et al. 2010, Kalkan, Olova et al. 2017). As *Hox* genes are highly enriched in SSPCs (**Fig. 1A**), we hypothesize that they maintain skeletal progenitors in an undifferentiated state. In support of this, in the adult zeugopod, *Hoxa11* is expressed in skeletal stem cells marked by, PDGFRα and CD51, and is absent from differentiated Osterix^+^ osteoblasts, Sox9^+^ chondrocytes, and Perilipin^+^ adipocytes (Rux, Song et al. 2016). However, comparatively little is known about *Hox* expression dynamics in the intermediate steps as cells transition from stem cells and start the differentiation process. If *Hoxa10* maintains cells in a primitive stem cell state, then one would expect its expression to be promptly shutdown when cells are challenged to differentiate. As PSCs are extremely rare, to investigate this we isolated PP1 early progenitor cells from the tibia by fluorescence activated cell sorting (FACS) and analyzed dynamic gene expression changes over the first 18 hours of osteogenic or adipogenic differentiation. Under both conditions, *Hoxa10* was rapidly downregulated by 2 hours of differentiation, even before known stem cell-associated genes (such as *Pdgfrα* and *CyclinD1*) were downregulated and lineage markers (like *FosB* and *Runx2*) were upregulated (**Fig. 2E, F**). These data suggest that *Hoxa10* must be downregulated for differentiation to proceed and underpins its potential role as a stem cell maintenance factor.

### Inhibition of Hox genes in stem and progenitor cells triggers a loss of skeletal stem cells and periosteal stemness properties

We hypothesize that *Hox* expression functions in skeletal stem cells, and not in more committed cells, to maintain the stem cell pool. In knockdown/knockout experiments, the role of individual *Hox* genes is frequently masked by the functional redundancy of other family members (Carpenter, Goddard et al. 1993, Mark, Lufkin et al. 1993, Studer, Lumsden et al. 1996, Gavalas, Studer et al. 1998, Rossel and Capecchi 1999, McNulty, Peres et al. 2005, Rux, Song et al. 2016, Song, Pineault et al. 2020). To test the maintenance role of *Hox* in the context of *in vivo* bone regeneration and to thwart potential redundancy of *Hox* genes, we utilized a conditional allele of the ∼100kb span of the *HoxA* cluster (Kmita, Tarchini et al. 2005) and paired it to three separate Cre drivers, *Pdgfrα^CreERT^, Osx^CreERT^,* and *Col1a1^CreERT^* that would target the *HoxA* family in increasingly more committed skeletal cells. Transgenic mice were given tamoxifen starting a week before tibial injury using a standardized tibial monocortical defect model, and EdU was also administered to the mice one day before sacrificing them to test proliferation capacity (**Fig. 3A**). Compared to control animals, knocking out the *HoxA* cluster in the *Pdgfrα^+^* stem and progenitor domain (Morikawa, Mabuchi et al. 2009, Pinho, Lacombe et al. 2013, Ambrosi, Longaker et al. 2019) during tibial injury led to a large reduction of 6C3^−^CD90^−^CD51^+^CD200^+^CD105^−^ skeletal stem cells, a concomitant increase in the more committed 6C3^−^CD90^−^CD51^+^CD200^−^CD105^−^ pre-Bone/Chondro/Stromal progenitors (pre-BCSPs), and a significant loss of proliferative capacity when probing cells harvested from the injury site at 3 days post injury by flow cytometry (**Fig. 3B**). This suggests that there may be a cell state shift as cells lose stemness and accumulate in this more committed cell state. The deletion of *HoxA* in more committed *Osx^+^* pre-osteoblasts (Mizoguchi, Pinho et al. 2014, Mabuchi, Okawara et al. 2021) resulted in a decrease in the pre-BCSP population without any change in proliferative activity (**Fig. 3C**). In contrast, *HoxA* deletion in *Col1a1^+^* mature osteoblasts (Kalajzic, Kalajzic et al. 2002) had no effect on skeletal stem and progenitor populations or proliferative capacity (**Fig. 3D**). Altogether, these results indicate that *Hox* genes are essential to the maintenance of skeletal stemness in the most primitive lineages present during skeletal regeneration.

**Figure 3.**
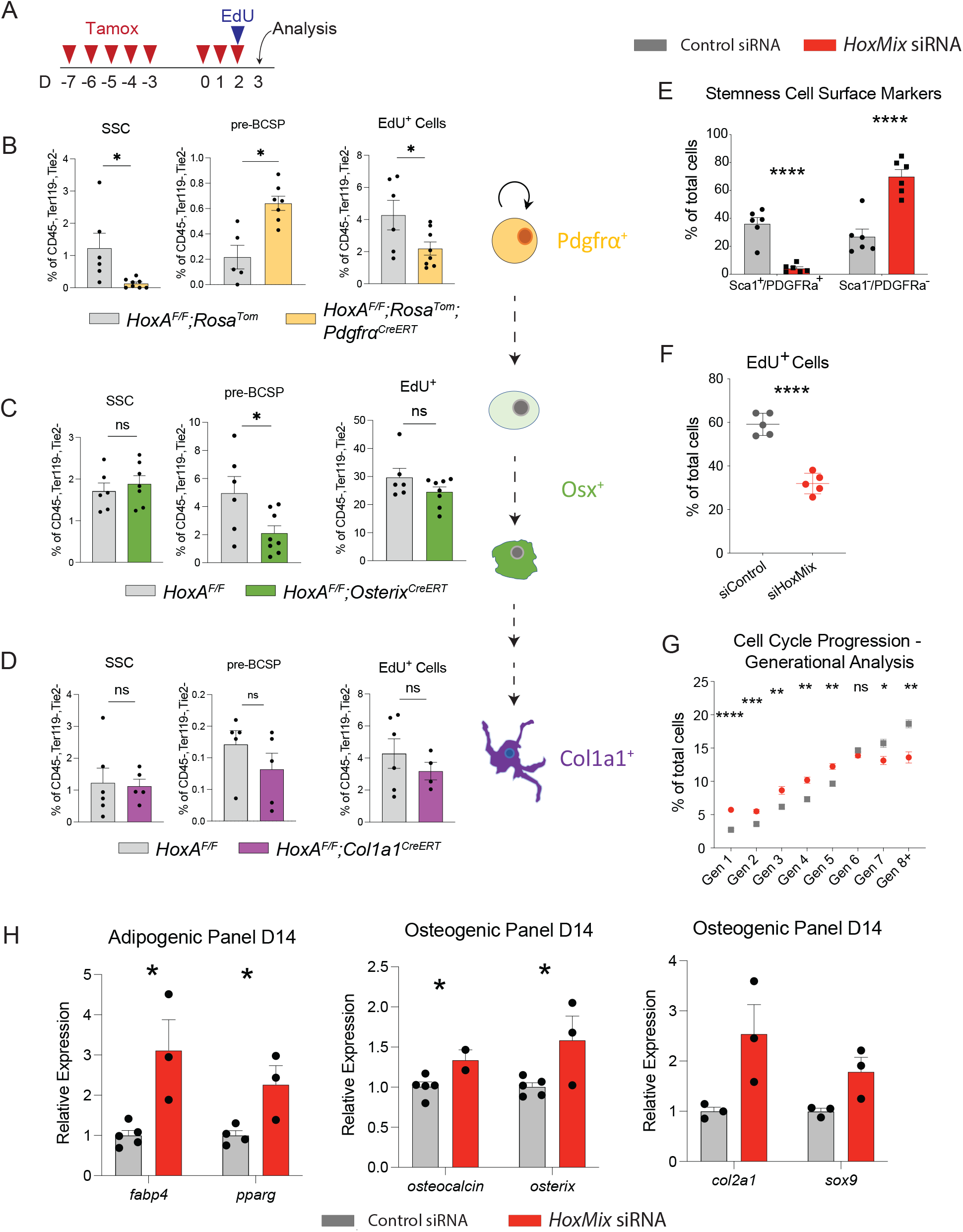
Inhibition of Hox genes in stem and progenitor cells triggers a loss of skeletal stem cells and periosteal stemness properties. (**A**) Scheme of tamoxifen dosing protocol (2mg/day) and EdU administration. (**B**-**D**) Flow cytometry revealed the percentage of 6C3^−^CD90^−^CD51^+^CD200^+^CD105^−^ skeletal stem cells, 6C3^−^CD90^−^ CD51^+^CD200^−^CD105^−^ pre-Bone/Chondro/Stromal progenitors (pre-BCSPs), and EdU^+^ proliferative cells in the nonhematopoietic compartment of *Pdgfr!^CreERT^;HoxA^flox/flox^* (**B**), *Osterix^CreERT^;HoxA^flox/flox^* (**C**), and *Col1a1^CreERT^;HoxA^flox/flox^* (**D**) and *HoxA^flox/flox^* control mice. (**E**-**F**) Simultaneous knockdown of *Hoxa10*, *Hoxa11*, *Hoxd10*, *Hoxd11*, and *Hoxc10* (*HoxMix*) was used to test the extent of stem cell potency in Hox-deficient tbial PSPCs. (**E**) After 7 days of control and *HoxMix* siRNA, PSPCs were analyzed for stemness-associated cell surface marker expression using flow cytometry. *n* = 3 each condition. (**F**) PSPCs were pulsed with EdU for 15 hours following HoxMix and nontargeting control siRNA knockdown; the amount of EdU-positive cells was then measured by flow cytometry. *n* = 5. (**G**) siControl and si*HoxMix* tibial PSPCs were also treated with Cell Trace^TM^ and subjected to flow cytometry to categorize cells by generation after six days of incubation. *n* =5 (control), *n* =7 (*HoxMix*). (**H**) Relative expression of adipogenic, osteogenic, and chondrogenic genes in tibial PSPCs serially transfected with control and *HoxMix* siRNA over the course of 14 days – measured by qRT-PCR. *n* = 5 (control), *n* = 3 (*HoxMix*). *ns* = not significant, \**p* < 0.05, \*\**p* < 0.01, \*\*\**p* < 0.001, \*\*\*\**p* < 0.0001. Two tailed Student’s t-test. Error bars are SEM.

To directly examine the role of *Hox* genes in periosteal stem and progenitor cells (PSPCs), we utilized an *si*RNA strategy to moderate their expression in isolated and cultured tibial periosteal cells, which are highly enriched for PSPCs (**Supplementary Fig. 1C, third panel**). To minimize potential redundancy between *Hox* genes highly expressed in these cells, we employed a mix of multiple siRNAs targeting *Hoxa10* and *11*, and *Hoxd10* and *11*, and *Hoxc10* (*HoxMix*), the most highly expressed *Hox* genes of the tibia periosteum. This strategy resulted in a significant reduction of the targeted *Hox* genes (**Supplementary Fig. 2A**). *Hoxa2*, a more proximal *Hox* gene, was not affected by the *HoxMix* siRNAs, confirming the specificity of this approach (**Supplementary Fig. 2A**). After seven days, we then assessed the impact of *Hox* knockdown on the skeletal stem cell state by analyzing the expression of the conventional skeletal stem cell markers SCA1 and PDGFRα (Houlihan, Mabuchi et al. 2012, Rux, Song et al. 2016) by flow cytometry. We found that cell cultures treated with *HoxMix* siRNAs had a significantly smaller proportion of cells that expressed both SCA1 and PDGFRα, while the size of the SCA1^−^/PDGFRα^−^ population increased, suggesting a shift from primitive undifferentiated to more mature, specified cells after *Hox* knockdown or priming towards more committed states (**Fig. 3E**).

Self-renewal and proliferation are hallmarks of stemness. The loss of *Hoxb4* and *Hoxa9* impairs proliferation and hinders the repopulating ability of hematopoietic stem cells (Bjornsson, Larsson et al. 2003, Brun, Bjornsson et al. 2004, Lawrence, Christensen et al. 2005). To determine whether a reduction in *Hox* genes also affects this functional property of skeletal stem cells, we performed an EdU incorporation assay on tibial periosteal cells after posterior *Hox* genes were downregulated using *HoxMix*. This demonstrated that proliferation and PSPC numbers decreased (**Fig. 3F**, **Supplementary Fig. 2D**), signifying the loss of stemness-associated characteristics. To investigate this further, we used CellTrace^TM^ to track cell cycle kinetics over multiple cell generations. In this assay cells were initially saturated with a fluorescent dye and, with each cell division, the dye is diluted in the daughter generation of cells. The amount of dye remaining in the cells after six days, as measured by flow cytometry, then revealed the generation number and cyling rate of cells assayed. Under control conditions, stem/progenitor cells predominantly comprise a higher-generation population of cells and are therefore high-cycling (**Supplementary Fig. 2B, C**). Following siRNA administration, generational analysis revealed that control cells continued to cycle and self-renew, with a large percentage being in the stemness-associated generation 7+, while *HoxMix* knockdown resulted in a reduction of cells in the higher generations (**Fig. 3G**), again supporting the hypothesis that *Hox* deficiency leads to a loss of stemness and lineage progression towards differentiation.

Downregulation or loss of genes associated with stemness can dismantle the self-renewal gene regulatory network and trigger aberrant spontaneous differentiation. For example, in the developing embryonic palate and other craniofacial skeletal elements, lack of *Hoxa2* induces an increase in differentiation and an upregulation of bone and cartilage markers (Kanzler, Kuschert et al. 1998, Dobreva, Chahrour et al. 2006, Iyyanar and Nazarali 2017). To probe the effect of *Hox* deficiency on tibial periosteal cell differentiation propensity, serial siRNA transfections were employed to suppress *Hox* for 14 days. Gene expression analysis confirmed the continued repression of *Hox* genes, and downregulation of the stem cell marker, *Pdgfrα*, in periosteal cells transfected with *HoxMix* (**Supplementary Fig. 2E, F**) and, in the absence of overt differentiation cues, these cells showed an increase in the expression of a cohort of osteogenic, adipogenic, and chondrogenic markers when compared to those treated with control siRNA (**Fig. 3H**). Together, these results demonstrate that, a reduction of *Hox* expression in skeletal stem and progenitor cells results in a loss of self-renewal capacity and an increase in spontaneous tri-lineage differentiation.

### Hoxa10 expression is sufficient to induce a skeletal stem cell state

The redundancy between *Hox* genes often obfuscates their function in loss-of-function experiments, hence the clearest indications for a role of HOX transcription factors in the regulation and maintenance of adult stem cells have come from overexpression studies (Sauvageau, Thorsteinsdottir et al. 1995, Lawrence, Sauvageau et al. 1996, Thorsteinsdottir, Sauvageau et al. 1997, Thorsteinsdottir, Mamo et al. 2002). As we found that *Hox* knockdown results in a cell fate shift towards a more mature skeletal cell phenotype, we asked whether, conversely, *Hoxa10* overexpression promotes stemness. We generated a lentiviral vector containing the protein-coding sequence of *Hoxa10* and that of *GFP* to mark infected cells (*LV-Hoxa10/GFP*), and a separate control vector containing only the *GFP* coding sequence (*LV-GFP*; **Fig. 4A**). The stable overexpression of *Hoxa10* was confirmed by qRT-PCR after infecting isolated periosteal stem and progenitor cells with *LV-Hoxa10/GFP* or *LV-GFP* (**Fig. 4B**). Compared to control PSPCs, which exhibited a large and broad shape morphology (**Fig. 4D**), *Hoxa10-*overexpressing PSPCs instead adopted a small, spindle shape morphology that is characteristic of typical mesenchymal stem cells ((Yang, Ogando et al. 2018); **Fig. 4D**). Moreover, seven days after infection, *Hoxa10* overexpressing periosteal cell cultures also had a significantly greater proportion of cells that expressed the stem cell markers PDGFRα, SCA1, and CD51 as assayed by flow cytometry (**Fig. 4C**). This is in accordance with the effect of *Hoxa10* on the hematopoietic system, where overexpression also increases the number of early hematopoietic progenitors (Magnusson, Brun et al. 2007). These data indicate that *Hoxa10* overexpression promotes a stem cell state and thus we next decided to probe how *Hoxa10* may regulate the capacity of tibial stem and progenitors to differentiate.

**Figure 4.**
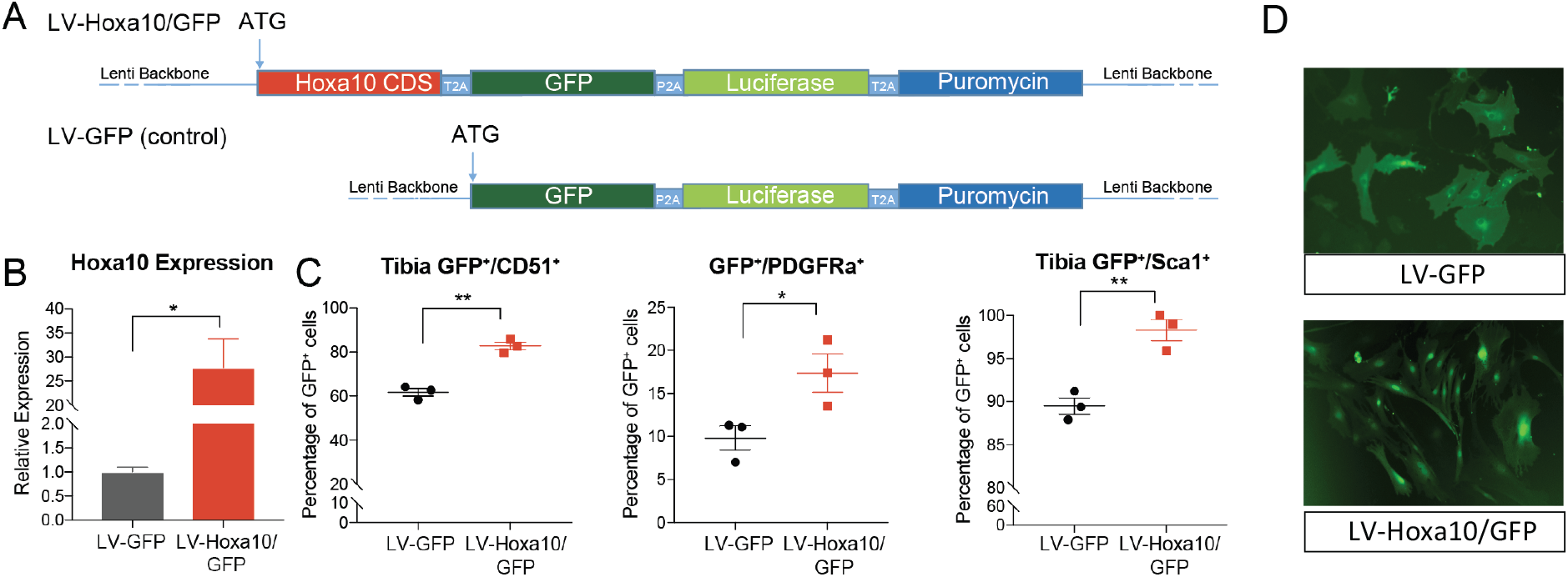
*Hoxa10* expression is sufficient to promote skeletal stem cell potency. (**A**) Schematic of lentiviral constructs used to transduce PSPCs. (**B**) qRT-PCR was used to reveal the relative expression of *Hoxa10* in control (LV-GFP) and *Hoxa10*-overexpressing (LV-Hoxa10/GFP) tibial PSPCs 11 days after infection. *n* = 3. (**C**) LV-GFP- and LV-Hoxa10/GFP-transduced tibial PSPCs were subjected to flow cytometry after a 7-day incubation to reveal the balance of infected GFP^+^ cells exhibiting the skeletal stem cell surface markers CD51, PDGFR*α*, and SCA1. *n* =3. (**D**) GFP fluorescent images demonstrating both the stable expression of the GFP marker and cell morphological differences after 7 days in LV-GFP- and LV-Hoxa10/GFP-transduced tibial PSPCs. \**p* < 0.05, \*\**p* < 0.01. Two tailed Student’s t-test. Error bars are SEM.

### Hoxa10-overexpressing periosteal stem and progenitor cells display deficient osteodifferentiation

Next, we investigated the effect of *Hoxa10* overexpression on the regulation of PSPC differentiation. Enforced expression of *Hox* genes blocks differentiation in numerous tissues and model organisms including *Hoxb4* and *Hoxa9* in lymphomyeloid differentiation (Owens and Hawley 2002, Schiedlmeier, Klump et al. 2003), *Hoxa5* during erythropoiesis (Crooks, Fuller et al. 1999, Fuller, McAdara et al. 1999), and *Hoxa2* during murine bone and cartilage development (Kanzler, Kuschert et al. 1998, Creuzet, Couly et al. 2002). The overexpression of *Hoxa10* inhibits commitment of early hematopoietic progenitors to the lymphomyeloid and erythroid lineages (Thorsteinsdottir, Sauvageau et al. 1997, Buske, Feuring-Buske et al. 2001, Taghon, Stolz et al. 2002, Magnusson, Brun et al. 2007) and blocks cardiac differentiation in cardiovascular progenitors (Behrens, Iacovino et al. 2013). This prompted us to ask whether *Hoxa10* plays a similar role in skeletal stem and progenitor cells of the tibia. To do so, we challenged *Hoxa10*-overexpressing cells to differentiate into the osteo-lineage (**Fig. 5A**). PSPCs that were infected with *LV-Hoxa10/GFP* and exposed to osteoinduction media for 14 days showed a more than 33% increase in the proportion of periosteal progenitors compared to those infected with the control virus, as measured by flow cytometry (**Fig. 5B**). *Hoxa10*-overexpressing cells also displayed lower expression of a suite of osteogenic genes (*Osterix*, *Osteocalcin*, and *Runx2)* after osteoinduction relative to control cells (**Fig. 5C**). Our findings suggest that *Hoxa10* overexpression either maintains cells in the stem cell-like state, while *LV-GFP* control cells begin to spontaneously differentiate or dysregulate – or *Hoxa10*-overexpressing cells are reprogrammed into a more stem cell-like state or a combination of both. We next investigated this proposition by probing the reprogramming abilities of *Hox* gene expression.

**Figure 5.**
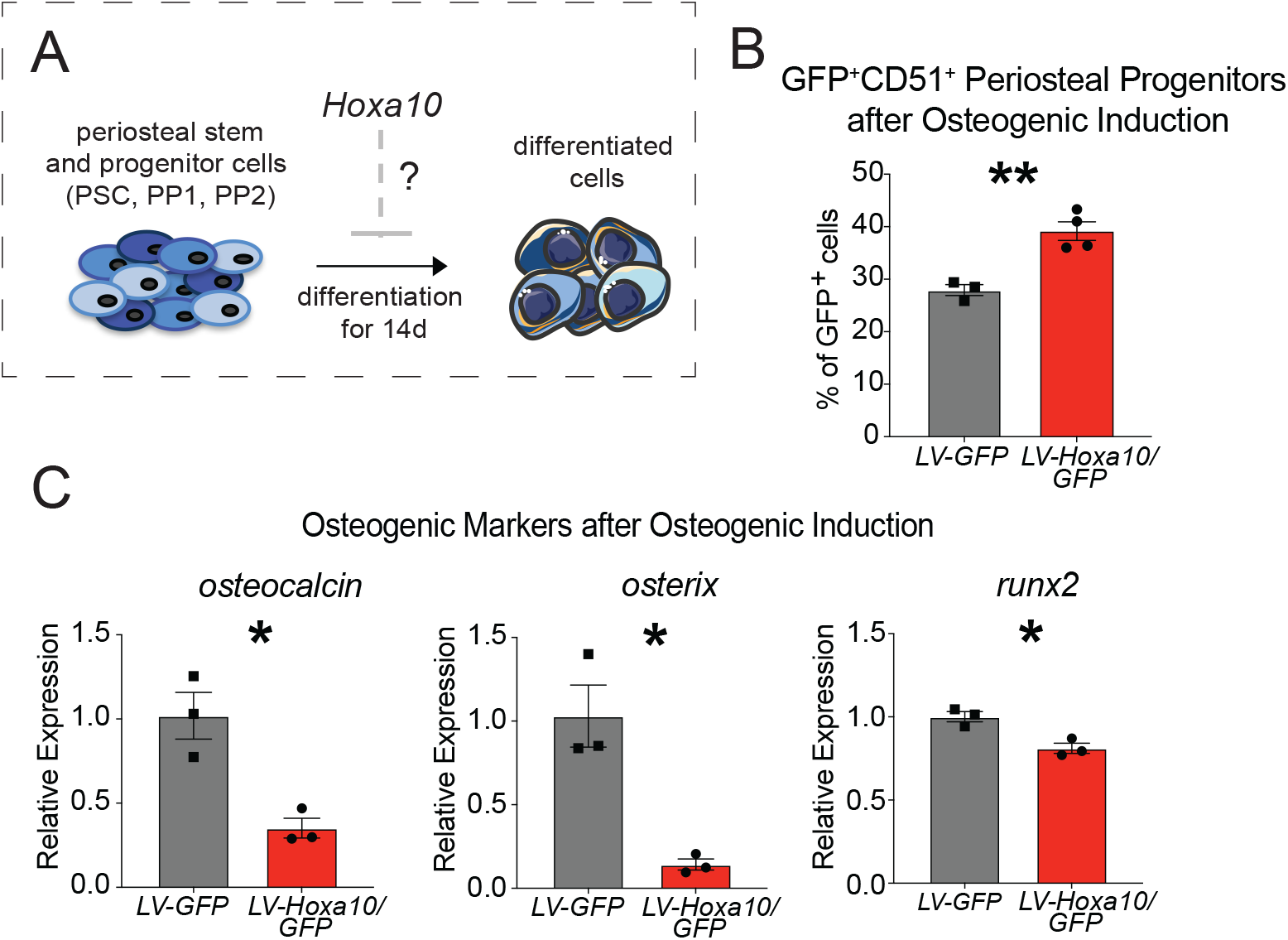
Hoxa10-overexpressing periosteal stem and progenitor cells display deficient osteodifferentiation. (**A**) Schematic of *in vitro* experiments performed in (**B**, **C**) to investigate whether *Hoxa10* overexpression can inhibit the differentiation of PSPCs after 14 days of osteogenic induction. (**B**) The balance of nonhematopoietic 6C3^−^CD90^−^ CD51^+^GFP^+^ PSPCs as a percentage of total infected cells (GFP^+^ cells) after transduction with *LV-GFP* or *LV-Hoxa10/GFP* and 14 days of osteoinduction media as measured by qRT-PCR. *n* = 3, *LV-GFP*; *n* = 4, *LV-Hoxa10/GFP* (**C**) Relative expression of osteogenic genes in *LV-GFP* or *LV-Hoxa10/GFP-*infected tibial periosteal stem and progenitors cells after a 14-day course of osteoinduction media. *n* = 3 each condition. \**p* < 0.05, \*\**p* < 0.01. Two tailed Student’s t-test. Error bars are SEM.

### Hoxa10 overexpression mediates reprogramming of PP1 cells into PSCs

Several Hox family members (including their Drosophila orthologues) function as pioneer factors, demonstrating a strong preference to bind inaccessible chromatin and, by doing so, increase accessibility (Beh, El-Sharnouby et al. 2016, Bulajić 2019). Cellular reprogramming is associated with an opening of chromatin (Gaspar-Maia, Alajem et al. 2011, Ugarte, Sousae et al. 2015) and we previously showed that *Hox*-positive periosteal cells have more open chromatin and are more stem-like than periosteal cells derived from *Hox*-negative tissue (Bradaschia-Correa, Leclerc et al. 2019). This, along with the increase in stem cell frequency described above, led us to postulate whether *Hox* expression can drive skeletal progenitors to a more primitive state.

To investigate this, we sorted each periosteal stem and progenitor population (PSC, PP1, and PP2), and separately transduced them with either *LV-Hoxa10/GFP* or *LV-GFP*, and reassessed the lineage hierarchy to see if some cells could convert to a more primitive state (**Fig. 6A**). After 7 days of incubation, FACS analysis revealed that the sorted PSC population largely shifted toward the PP1 cell state, suggesting the PSCs rapidly differentiate to this more committed state in these culture conditions. However, no difference in cell populations within the PSPC lineage hierarchy was observed between PSCs transduced with *LV-Hoxa10/GFP* or *LV-GFP* (**Fig. 6C**). Strikingly, in sorted PP1 cells that were *LV-Hoxa10/GFP-* infected, FACS analysis revealed an increase in the more uncommitted stem cell compartment at the expense of the PP1 population after 7 days (16% converted cells); this proportional increase was limited in the *LV-GFP*-infected PP1 cells (5% converted cells)(**Fig. 6B, D, and Supplementary Fig. 3A**). This limited amount of stem cells in the control-infected PP1 cells may reflect a basal level of stochastic sampling of neighboring cellular states, as has been described in many other purified cell populations (Gupta, Fillmore et al. 2011, Wang, Quan et al. 2014). In contrast, PP2 cells showed no difference in the relative abundance of the different cell populations when overexpressing *Hoxa10* (**Fig. 6E**). Multiple independent iterations of this experiment confirmed these results, in which *Hoxa10*-overexpressing PP1 cells led to a greater than two-fold increase in the proportion of uncommitted periosteal stem cells among cells in the PSPC compartment (**Fig. 6F; Table 1**) and around a three-fold increase in the frequency of PSCs among total *Hoxa10*-overexpressing cells (**Fig. 6G**).

**Figure 6.**
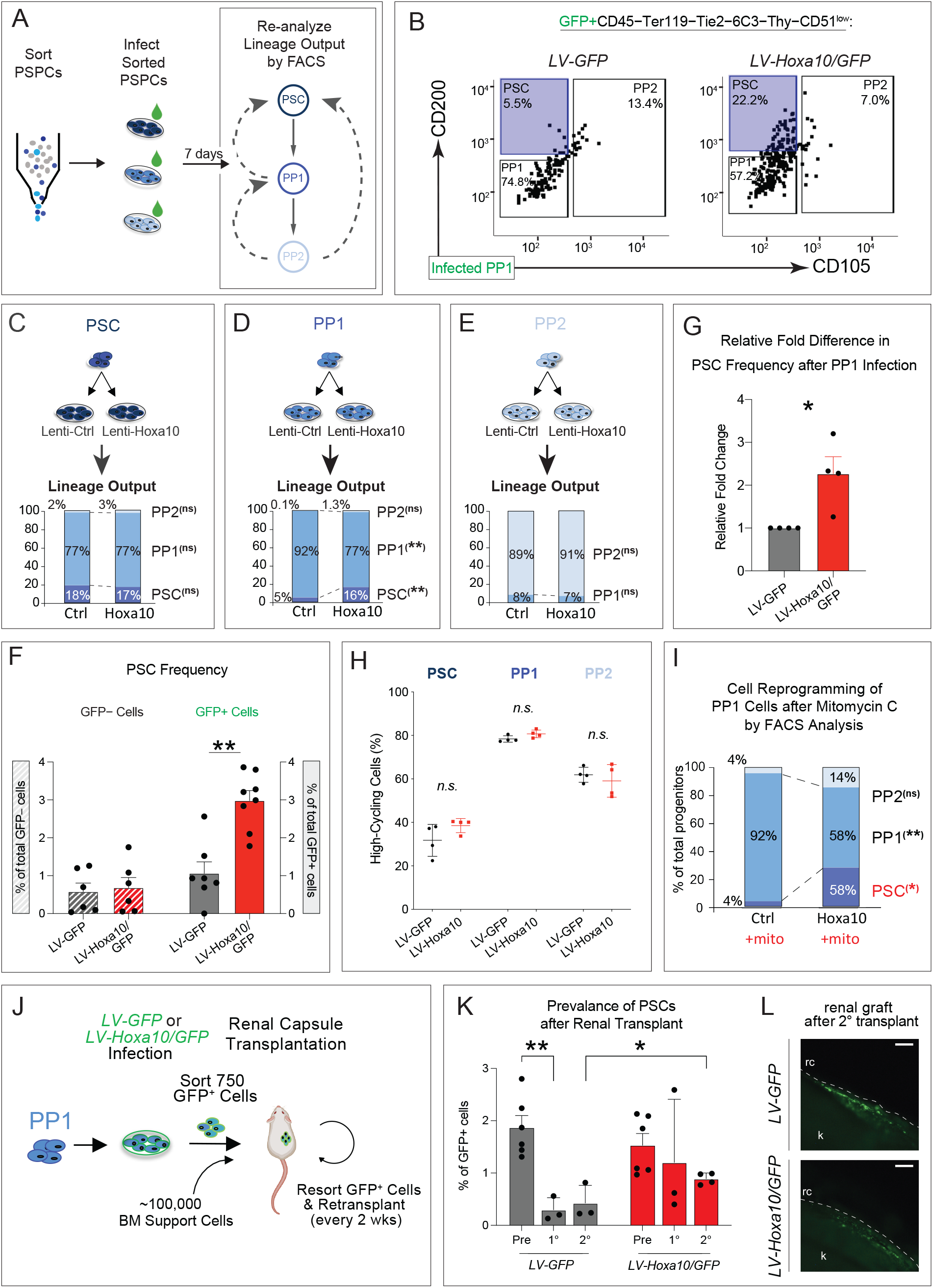
Hoxa10 overexpression mediates reprogramming of adult tibial PP1 cells into PSCs. (**A**) Experimental plan to test the reprogramming abilities of *Hoxa10.* Tibial PSCs, PP1 and PP2 cells were separately isolated by FACS. Each cell population was infected with *LV-GFP* or *LV-Hoxa10/GFP* and analyzed by flow cytometry after 7 days of incubation. (**B**) Representative FACS plots of the PSPC lineage output 7 days after transduction of PP1 cells with *LV-GFP* or *LV-Hoxa10/GFP*. Only infected (GFP^+^) cells were evaluated. (**C**-**E**) Flow cytometric analysis of the distribution of GFP^+^ PSCs, PP1, and PP2 cells within the CD51^+^ stem and progenitor cell compartment 7 days after *LV-GFP* or *LV-Hoxa10/GFP* infection of PSCs (**C**), PP1 (**D**), and PP2 (**E**) cells. *n* = 3 (PSC), *n* = 3 (PP1), *n* = 5 (PP2). (**F**) The relative fold change in GFP^+^ PSCs within the PSPC compartment after transduction of PP1 cells with *LV-GFP* or *LV-Hoxa10/GFP. n* = 4 separate experiments. (**G**) The frequency of PSCs among total cells 7 days after the infection of PP1 cells with *LV-GFP* or *LV-Hoxa10/GFP*. Uninfected (GFP^−^) and infected (GFP^+^) are shown separately. (**H**) *LV-GFP*- or *LV-Hoxa10/GFP*-infected tibial PSPCs were also treated with Cell Trace^TM^ and subjected to flow cytometry to categorize cells as high- or low-cycling after six days of incubation. Gating strategy is presented in **Supplementary Fig. 3A.** *n* = 4 each condition. (**I**) Flow cytometry revealed the lineage hierarchy of tibial PP1 cells 7 days after 10ug/mL mitomycin C treatment and infection with *LV-GFP*- or *LV-Hoxa10/GFP. n* = 3. (**J**) Experimental plan to carry out serial transplants of reprogrammed periosteal cells under the renal capsule to test self-renewal capacity. (**K**) PP1 cells were first transduced with either *LV-GFP* or *LV-Hoxa10/GFP* and the prevalence of GFP-labelled PSCs was then assessed by flow cytometry before (Pre) and after each round of transplantation. *n* = 6, Pre-*LV-GFP* and Pre-*LV-Hoxa10/GFP*; *n* = 3, 1^○^-*LV-GFP* and 1^○^-*LV-Hoxa10/GFP*; *n* = 3, 2^○^-*LV-GFP*; *n* = 4, 2^○^-*LV-Hoxa10/GFP.* (**L**) Representative fluorescent images of renal capsule grafts derived from *LV-GFP* or *LV-Hoxa10/GFP*-infected periosteal cells after two rounds of transplantation (rc = renal capsule, k = kidney). Scale bars, 200 μm. (**B**-**E**, **G**) Complete results and statistics are provided in **Table 1.** *n.s.* = not significant, \**p* < 0.05, \*\**p* < 0.01. Two tailed Student’s t-test. Error bars are SEM.

**Table 1.**
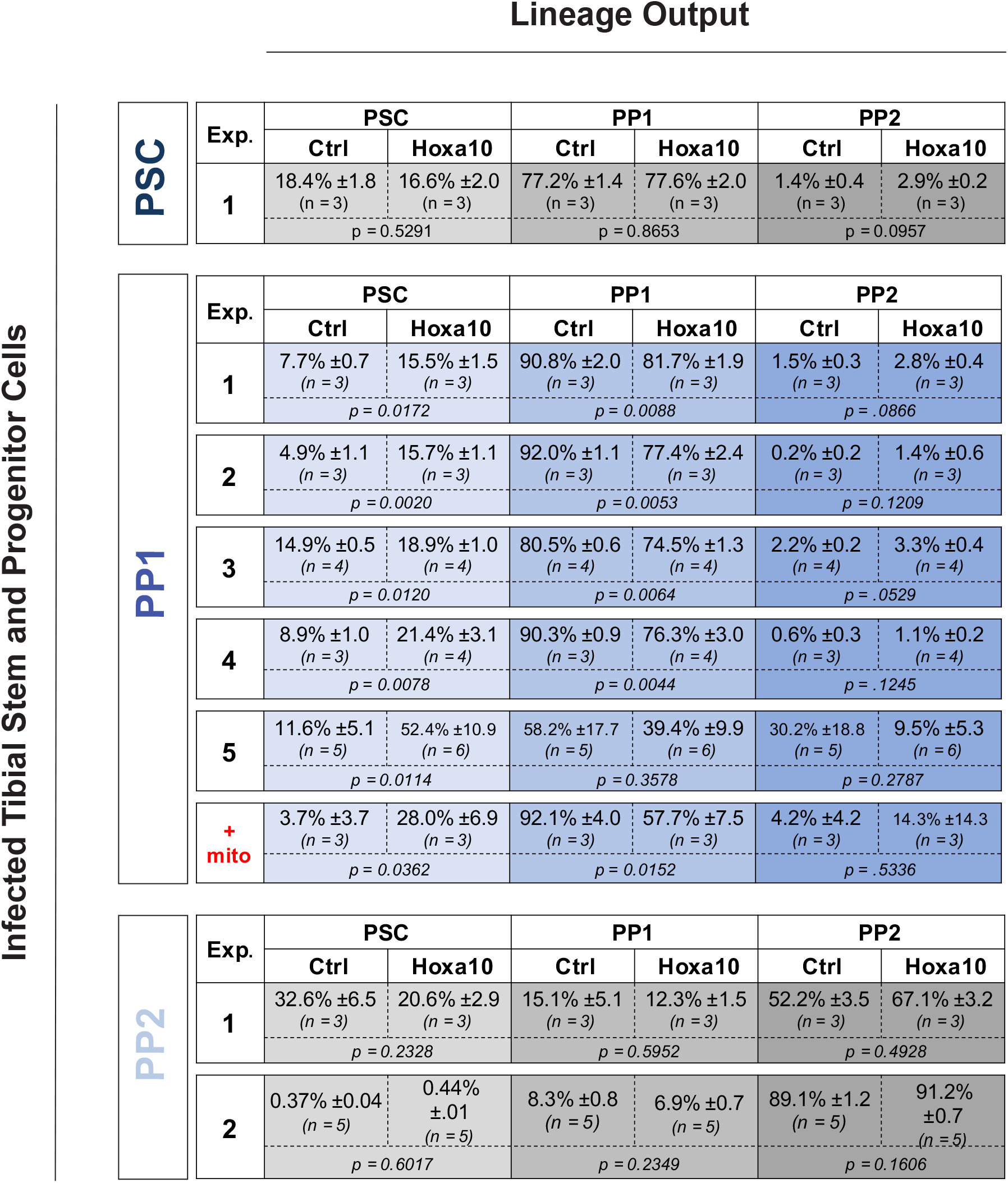
The lineage output after *Hoxa10* overexpression in adult tibial PSPCs. The lineage hierarchy of sorted PSCs, PP1 or PP2 cells was evaluated by flow cytometry seven days after transduction with *LV-GFP* (Ctrl) or *LV-Hoxa10* (Hoxa10). Only infected GFP^+^ cells were assessed. Multiple separate experiments (Exp.) are shown.

To examine the possibility that *Hoxa10* overexpression merely triggers proliferation of the more uncommitted periosteal cells, we employed CellTrace^TM^ to assess the cycling rate of each stem and progenitor compartment infected with *LV-Hoxa10/GFP* or *LV-GFP*. In these experiments, *Hoxa10* overexpression did not increase the proliferative rate of PSPCs (**Fig. 6H, Supplementary Fig. 3B**). This contrasted with the *Hox* knockout and *si*RNA knockdown experiments in which *Hox* expression had a significant influence on proliferative capacity (**Fig. 3**). One explanation may be that there is a maximum proliferation rate that PSPC populations can reach. Consistent with this, studies have shown that PSPCs are already highly proliferative, displaying the highest proliferative capacity when compared to other mesenchymal progenitor populations derived from different tissues (Sakaguchi, Sekiya et al. 2005, Yoshimura, Muneta et al. 2007, van Gastel, Torrekens et al. 2012, Duchamp de Lageneste, Julien et al. 2018). We further confirmed the insignificance of proliferation for PSC conversion by inhibiting proliferation in PP1 *Hoxa10*-overexpressing cells via mitomycin C treatment. After confirming effective inhibition of proliferation (**Supplementary Fig. 3C**), we again examined the lineage hierarchy and observed a cell fate switch from the progenitor to the more primitive stem cell (**Fig. 6I**). Interestingly, the magnitude of the conversion was much larger than in the case of PP1 cells with unaltered proliferative capacity. Mitomycin C is known to arrest cells in the G1 phase of the cell cycle (Kang, Chung et al. 2001) and multiple studies have connected cell fate decisions to a prolonged G1 (Sela, Molotski et al. 2012, Tapias, Zhou et al. 2014). It may be that PP1 cells in cell cycle arrest create a more permissive window for cell reprogramming factors to change their cell state. Overall, these data highly suggest that the expansion of stem cells among total cells and within the PSPC compartment is due to progenitors reprogramming to a less committed state.

To further assess whether the more committed PP1 cells have genuinely been reprogrammed to a periosteal stem cell fate, we utilized published single-cell gene expression data (Debnath, Yallowitz et al. 2018) to identify new transcriptional markers of periosteal stem cells. We sorted cells *in silico* using the previously defined PSPC cell surface markers and uncovered several new factors that were additionally enriched in periosteal stem cells versus PP1 and PP2 cells (*Omd*, *Car3*, *Ucma*, and *Frzb* (**Supplementary Fig. 4A**). Gene expression analysis of PP1 cells transduced with *LV-Hoxa10/GFP* by qRT-PCR revealed a higher expression of these newly identified periosteal stem cell marker genes relative to those infected with *LV-GFP* (**Supplementary Fig. 4B**), indicating a more comprehensive shift towards the distinct stem cell transcriptome as cells are reprogrammed. Stem cells are defined by their capacity to self-renew and differentiate. PSCs are at the top of the periosteal skeletal stem and progenitor lineage hierarchy and thus possess a greater ability to self-renew during serial transplantation assays relative to PP1 and PP2 cells (Debnath, Yallowitz et al. 2018). To investigate whether PP1 *Hoxa10*-overexpressing cells are functionally reprogrammed to a stem cell state, we interrogated their competence to self-renew using a serial transplantation assay in which PP1 cells are first infected with either *LV-Hoxa10/GFP* or *LV-GFP*, and 300 to 750 GFP^+^ cells are isolated and transplanted (along with 100,000 bone marrow support cells) underneath the renal capsule for two weeks – an environment that promotes differentiation of mesenchymal stem cells (Debnath, Yallowitz et al. 2018, Ambrosi, Marecic et al. 2021). GFP^+^ cells were then re-sorted and re-transplanted for a further two weeks and the persistence of GFP^+^ PSCs (reprogrammed from PP1 cells) was determined following each transplantation (**Fig. 6J**). Indeed, flow cytometry of post-transplantation cells revealed that GFP+ PSCs derived from control *LV-GFP-*infected PP1 cells were largely lost over time when compared to the pre-transplantation frequency. In contrast, PSCs derived from *LV-Hoxa10/GFP*-infected PP1 cells were maintained to a larger degree after multiple rounds of transplantation, indicating a greater self-renewal capacity (**Fig. 6K, L**). With the capacity of *Hoxa10* to shift periosteal progenitors of the tibia towards less committed state that possesses greater self-renewal capacity, these results highlight a master role for *Hox* genes in the regulation of skeletal stem cells.

### The regional specificity of Hox function is maintained in the adult skeleton

Previous studies, including findings from our own lab, showed that *Hox* gene expression is maintained in adult skeletal elements other than the tibia and that the expression patterns roughly mirror the regionally restricted expression during embryogenesis (**Fig. 7I**) (Ackema and Charite 2008, Leucht, Kim et al. 2008, Rux, Song et al. 2016, Rux and Wellik 2017, Bradaschia-Correa, Leclerc et al. 2019, Song, Pineault et al. 2020). Thus, as in development, the adult “Hox code” may endow stem cells from different anatomical locations with the specific functional properties needed to successfully regenerate the tissue in which they reside. While periosteal cells derived from distinct skeletal regions exhibit functional differences that are correlated with differential *Hox* expression (Leucht, Kim et al. 2008, Bradaschia-Correa, Leclerc et al. 2019), the regional specificity of *Hox* function in the adult has not been definitively established. To investigate this, we asked whether *Hoxa10* functions in a universal manner in stem and progenitor cells from any part of the skeleton or whether its stem cell maintenance function is limited to the tibia, with, perhaps, other *Hox* genes filling that role in other regions. First, we isolated periosteal cells from various skeletal elements, including the pelvis, the thoracic vertebrae 5 through 8 of the spine, the radius/ulna, and anterior ribs 1 through 4 (**Fig. 7A**), and subjected them to gene expression analysis using the Nanostring nCounter^®^ technology to uncover the *Hox* expression profiles of all 39 *Hox* genes among these four tissues. We found that, as in the tibia, *Hoxa10* is the most highly expressed family member in periosteal cells of the pelvis and radius/ulna (**Fig. 7A, E**; **Supplemantary Fig. 6**). In the adult spine^T5-8^ periosteum, *Hoxb8* is the highest expressed (**Fig. 7C; Supplementary Fig. 6**), reflecting the developmental expression profile as this section of the vertebral column is configured by the Hox8 group (Favier and Dolle 1997, van den Akker, Reijnen et al. 1999). Finally, *Hoxa5* is the most highly expressed *Hox* gene in the anterior ribs^1-4^ (**Fig. 7G**).

**Figure 7.**
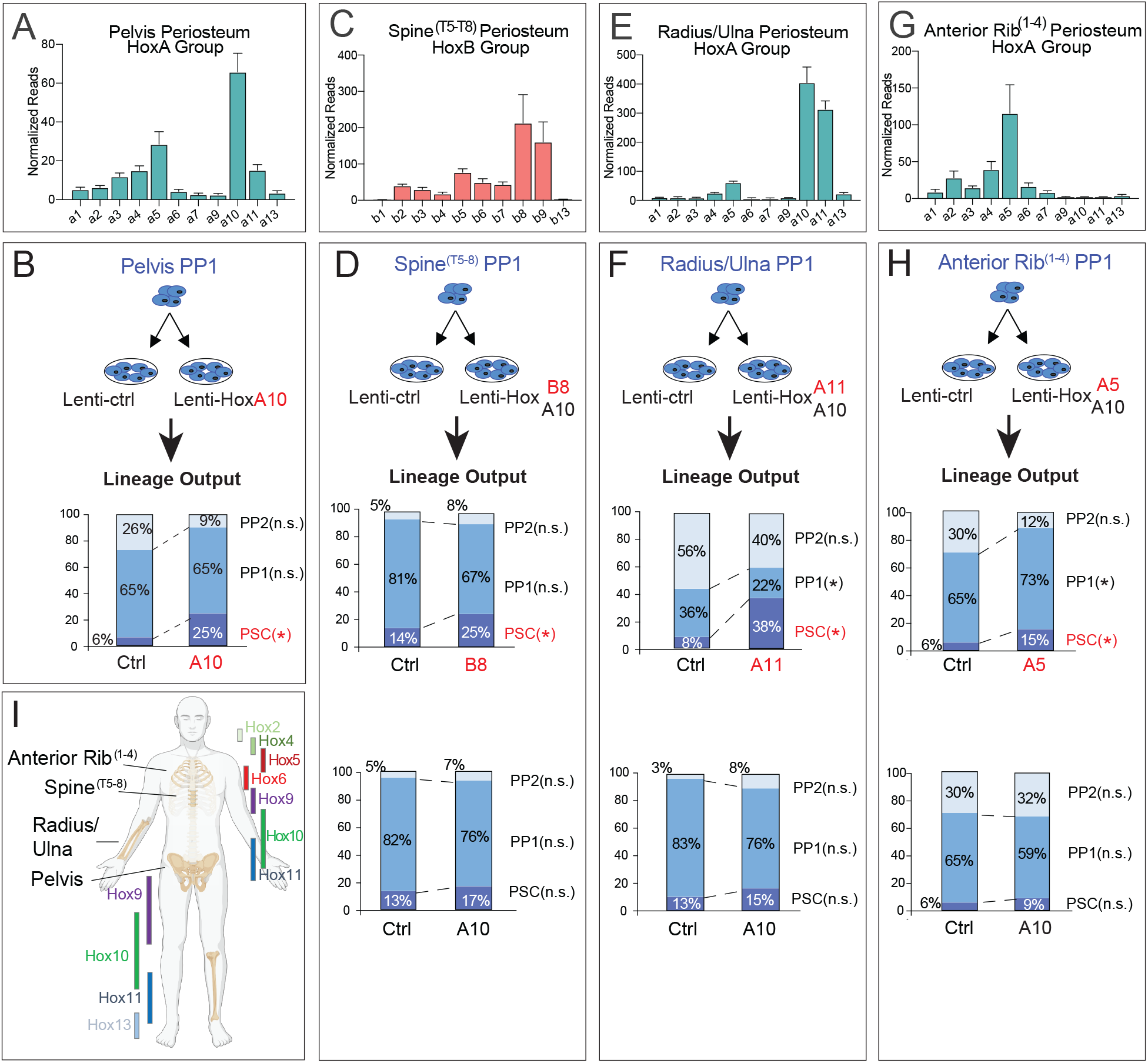
The regional specificity of *Hox* function is maintained in the adult skeleton. (**A, C, E, G**) The expression profile of the Hox cluster containing the highest expressed Hox gene in periosteal cells of the pelvis (**A**), spine^T5-T8^ (**C**), radius/ulna (**E**), and anterior rib^1-4^ (**G**). *n* = 4 mice for each skeletal element. Full Hox expression data in **Supplementary Fig. 6**. (**B**, **D**, **F**, **H**) The lineage output of stem and progenitors 7 days after infecting PP1 cells deriving from the pelvis (**B**), spine^T5-T8^ (**D**), radius/ulna (**F**), and anterior rib^1-4^ (**H**) with either *LV-Hoxa10/GFP*, *LV-Hoxb8/GFP*, *LV-Hoxa11/GFP, or LV-Hoxa5/GFP*, respectively – and with *LV-GFP* (Ctrl) and *LV-Hoxa10/GFP* serving as a control. *n* = 5, pelvis (Ctrl and A10); *n* = 4, spine (Ctrl and B8); *n* = 3, spine (Ctrl and A10); *n* = 4 and *n* = 5, radius/ulna (Ctrl and A11, respectively); *n* = 5, radius/ulna (Ctrl and A10); *n* =9, *n* = 8, and *n* = 9, anterior rib (Ctrl, A5, and A10, respectively). Full lineage output data and statistics are provided in **Table 2**. (**I**) Diagram of skeletal elements investigated along with the proposed regional restriction Hox expression in adult skeletal tissues (adapted from *Rux and Wellik, 2017*). *n.s.* = not significant, \**p* < 0.05. Two tailed Student’s t-test. Error bars are SEM.

**Table 2.** The lineage output after *Hox* overexpression in adult PSPCs from various anatomical regions. The lineage hierarchy of sorted PSCs, PP1 or PP2 cells was evaluated by flow cytometry seven days after transduction with *LV-GFP* (Ctrl), *LV-Hoxa10* (Hoxa10), or a lentivirus containing the Hox gene shown to have the highest expression in each corresponding skeletal element. Only infected GFP^+^ cells were assessed. Multiple separate experiments are shown.

|  |  |  |  |  |  |  |  |
| --- | --- | --- | --- | --- | --- | --- | --- |
| Tibia PP1 | Hox | PSC |  | PP1 |  | PP2 |  |
|  |  | Ctrl | Hox | Ctrl | Hox | Ctrl | Hox |
|  | A5 | 11.6% ±5.1<br>(n = 5) | 15.8% ±3.9<br>(n = 6) | 58.2% ±17.7<br>(n = 5) | 72.4% ±4.7<br>(n = 6) | 30.2% ±18.8<br>(n = 5) | 11.8% ±1.4<br>(n = 6) |
|  |  | p = 0.5299 |  | p = 0.4185 |  | p = 0.3078 |  |
|  | A11 | 11.6% ±5.1<br>(n = 5) | 32.4% ±5.7<br>(n = 6) | 58.2% ±17.7<br>(n = 5) | 58.2% ±6.9<br>(n = 6) | 30.2% ±18.8<br>(n = 5) | 9.4% ±2.7<br>(n = 6) |
|  |  | p = 0.0260 |  | p = 0.9982 |  | p = 0.2573 |  |
| Spine<sup>T5-8</sup> PP1 | Hox | PSC |  | PP1 |  | PP2 |  |
|  |  | Ctrl | Hox | Ctrl | Hox | Ctrl | Hox |
|  | A10 | 12.9% ±1.7<br>(n = 3) | 16.7% ±2.7<br>(n = 3) | 79.0% ±4.1<br>(n = 3) | 75.7% ±1.6<br>(n = 3) | 4.2% ±2.1<br>(n = 3) | 6.6% ±0.4<br>(n = 3) |
|  |  | p = 0.2994 |  | p = 0.4950 |  | p = .3695 |  |
|  | B8 | 14.9% ±1.3<br>(n = 4) | 24.6% ±4.0<br>(n = 4) | 80.7% ±2.1<br>(n = 4) | 67.5% ±4.3<br>(n = 4) | 5.4% ±1.1<br>(n = 4) | 7.9% ±0.7<br>(n = 4) |
|  |  | p = 0.0435 |  | p = 0.0338 |  | p = 0.0964 |  |
|  | B8 | 12.3% ±3.3<br>(n = 4) | 31.8% ±5.5<br>(n = 4) | 60.2% ±8.2<br>(n = 4) | 48.9% ±7.1<br>(n = 4) | 25.5% ±10.9<br>(n = 4) | 19.4% ±1.6<br>(n = 4) |
|  |  | p = 0.0120 |  | p = 0.0064 |  | p = .0529 |  |
| Ant. Rib<sup>1-4</sup> PP1 | Hox | PSC |  | PP1 |  | PP2 |  |
|  |  | Ctrl | Hox | Ctrl | Hox | Ctrl | Hox |
|  | A10 | 10.2% ±0.5<br>(n = 6) | 12.5% ±2.7<br>(n = 6) | 80.4% ±2.3<br>(n = 6) | 83.4% ±3.3<br>(n = 6) | 9.4% ±2.1<br>(n = 6) | 3.1% ±1.4<br>(n = 6) |
|  |  | p = 0.0606 |  | p = 0.5727 |  | p = 0.3189 |  |
|  | A10 | 5.7% ±2.8<br>(n = 9) | 8.9% ±3.5<br>(n = 9) | 64.5% ±11.3<br>(n = 9) | 59.5% ±10.9<br>(n = 9) | 29.8% ±11.8<br>(n = 9) | 31.6% ±12.7<br>(n = 9) |
|  |  | p = 0.4824 |  | p = 0.7528 |  | p = 0.9168 |  |
|  | A5 | 5.7% ±2.8<br>(n = 9) | 15.4% ±2.5<br>(n = 8) | 64.5% ±11.3<br>(n = 9) | 73.1% ±4.0<br>(n = 8) | 29.8% ±11.8<br>(n = 9) | 11.6% ±2.5<br>(n = 8) |
|  |  | p = 0.0218 |  | p = 0.4875 |  | p = 0.1504 |  |
| R/U PP1 | Hox | PSC |  | PP1 |  | PP2 |  |
|  |  | Ctrl | Hox | Ctrl | Hox | Ctrl | Hox |
|  | A10 | 13.3% ±5.3<br>(n = 5) | 15.3% ±1.8<br>(n = 5) | 83.3% ±4.3<br>(n = 5) | 76.0% ±2.4<br>(n = 5) | 3.4% ±1.2<br>(n = 5) | 8.3% ±1.3<br>(n = 5) |
|  |  | p = 0.7350 |  | p = 0.1746 |  | p = 0.0253 |  |
|  | A11 | 8.3% ±5.3<br>(n = 5) | 37.5% ±1.1<br>(n = 4) | 37.8% ±8.3<br>(n = 5) | 22.5% ±3.4<br>(n = 4) | 56.0% ±2.4<br>(n = 5) | 40% ±11.2<br>(n = 4) |
|  |  | p = 0.01967 |  | p = 0.2481 |  | p = 0.222 |  |
| Pelvis PP1 | Hox | PSC |  | PP1 |  | PP2 |  |
|  |  | Ctrl | Hox | Ctrl | Hox | Ctrl | Hox |
|  | A11 | 6.1% ±1.8<br>(n = 5) | 24.6% ±4.2<br>(n = 5) | 65.2% ±14.7<br>(n = 5) | 65.5% ±3.6<br>(n = 5) | 26.4% ±12.8<br>(n = 5) | 9.6% ±1.8<br>(n = 5) |
|  |  | p = 0.7350 |  | p = 0.1746 |  | p = 0.0253 |  |

Periosteal progenitors were then isolated from each anatomical region and infected with either control, *Hoxa10*, or the Hox gene shown to be most highly expressed in that region. In the pelvis, spine^T5-8^, and anterior ribs^1-^4, PP1 cells were only reprogrammed to a PSC state when transduced with their correct regional-specific Hox gene (**Fig. 7; Supplementary Fig. 6; Table 2**). PP1 cells overexpressing a Hox gene from another region, for example, *Hoxa10* expression in periosteal cells from the spine or rib, did not enhance reprogramming relative to the control virus. Interestingly, in the radius/ulna towards a stem cell fate despite being the most expressed Hox gene in the periosteum of this region. *Hoxa11*, however, is also very highly expressed in this skeletal element and *Hoxa11* lentiviral overexpression in this region induced reprogramming of radius/ulna PP1 cells. It is noteworthy that in whole-body knockouts of *Hoxa10*, only minor skeletal defects in the radius and ulna are observed when compared to defects that arise in *Hoxa11* mutants (Small and Potter 1993, Davis, Witte et al. 1995, Favier, Rijli et al. 1996, Boulet and Capecchi 2002, Wellik and Capecchi 2003), suggesting that the function of periosteal cells in this region may be more influenced by *Hoxa11*.

As embryonic development progresses anterior-to-posterior (and proximal-to-distal in the limbs), each Hox gene cluster successively becomes more accessible and expressed, generating a nested pattern where more posterior tissues express a wider set of Hox genes than anterior ones (Tarchini and Duboule 2006, Papageorgiou 2012). This prompts the hypothesis that more posterior tissues may respond to anterior *Hox* gene expression. To test this, we infected tibial PP1 cells with *Hoxa5*-overexpressing or control lentivirus. *Hoxa5* overexpression in these posterior progenitor cells did not induce reprogramming (**Supplementary Fig. 7A**). Along with Hoxa10, Hoxa11 is also highly expressed in tibia periosteal cells and, when overexpressed, triggered tibia periosteal progenitor cells to revert to a more primitive state (**Supplementary Fig. 7B**). Altogether, these results reveal a model in which *Hox* genes function to maintain periosteal stemness and as reprogramming modulators in a regionally restrictive manner – and suggests that there is a limited set of ‘flanking’ Hox genes to which periosteal progenitors can respond. Importantly, this may have wider implications on the engraftment potential of future cell transplant therapies where tissue is often taken from one part of the body and transplanted into another.

## Discussion

Skeletal homeostasis and regeneration rely heavily on stem cells to replenish the tissue lost due to injury or wear and tear. To preserve this capacity, stem cell number has to be constantly maintained by cell division, one of the hallmarks of stemness. There is a relatively poor understanding of the molecular mechanisms that govern skeletal stem and progenitor cell maintenance and lineage progression during bone healing, and this presents one of the major hurdles to advancing cell-based therapies for treatment of bone fractures. This investigation adds to the growing appreciation that *Hox* genes have important maintenance functions in the adult skeleton. The experiments herein support a model in which high expression of *Hox* genes in the most uncommitted periosteal cell compartments confers greater proliferative ability, self-renewal capacity, and inhibits lineage progression towards more committed cell fates. They also demonstrate that this role of Hox can be exploited to shift periosteal progenitors with limited self renewal capacity (Debnath, Yallowitz et al. 2018) to a more primitive state, thus increasing their functional potential.

Previous reports have identified *Hoxa11* as the primary *Hox* gene that patterns the embryonic tibia and is a marker of the adult tibia (Wellik and Capecchi 2003, Rux, Song et al. 2016) Our work revealed that *Hoxa10*, not *Hoxa11*, is the highest expressed Hox gene in the tibial periosteum (**Supplementary Fig. 1**). Although the Hox11 group is regionally restricted to both the radius/ulna and the tibia/fibula in the adult (Rux, Song et al. 2016), only the radius and ulna fail to develop in Hoxa11/Hoxd11 double mutants. The tibia and fibula are only lightly affected (Davis, Witte et al. 1995, Wellik and Capecchi 2003), suggesting that these genes play a more substantial role in the forelimbs. As we identified *Hoxa10* as the most expressed Hox gene in the adult tibia periosteum, we hypothesized that this paralog may have a functional role in this skeletal element. To date, only whole embryonic (non-inducible) *Hox10* group knockouts have been studied. Both *Hoxa10*^−/−^ and *Hoxd10*^−/−^ mice display skeletal patterning alterations in the developing hindlimbs, with changes in these structures appearing with greater penetrance in *Hoxa10^−/−^* mice (Wahba, Hostikka et al. 2001).

While recent studies demonstrate that *Hoxa11* marks a primitive mesenchymal stem cell (MSC) in the periosteum and that *Hoxa11*-lineage marked cells are long-term contributors to MSCs throughout life (Rux, Song et al. 2016, Pineault, Song et al. 2019, Song, Pineault et al. 2020), we show for this first time that *Hox* genes play a stem cell maintenance function in the skeletal system. During differentiation of osteoblastic cell lines, HOXA10 drives the early expression of osteogenic genes through chromatin remodeling, and the *in vivo* conditional deletion of *Hoxa11* and *Hoxd11* in the *Hoxa11* domain leads to osteogenic differentiation defects (Hassan, Tare et al. 2007, Song, Pineault et al. 2020) suggesting a role for some Hox genes at later stages of cell fate commitment. Notably, using a *Hoxa11* knock-in reporter that simultaneously deletes *Hoxa11* coding sequence, *Song et al.* found that skeletal stem cells expressing the reporter are still present 10 months after the conditional deletion of *Hoxa11* and *Hoxd11 alleles* in 8-week-old forelimbs (Song, Pineault et al. 2020), presumably precluding a stem cell maintenance role for *Hox* genes. These studies focus on the *Hox11* paralogous group, however, and disregard the potential functional redundancy of other *Hox* groups expressed at high levels in the skeletal cells under study – as we observe in the tibia periosteum (**Fig. 2A, Supplementary Fig. 1**).

Recent loss-of-function studies in various tissues have demonstrated a stem cell maintenance role for *Hox* genes although phenotypes have usually been mild (Magli, Largman et al. 1997, Owens and Hawley 2002, Bjornsson, Larsson et al. 2003, Brun, Bjornsson et al. 2004, Iyyanar and Nazarali 2017). Additionally, *Rux et al.* have recently shown that skeletal stem cells harvested from mice in which both alleles of *Hoxd11* and one allele of *Hoxa11* are knocked out display tri-lineage differentiation dysfunction but do not lose their stemness marker profile or self-renewal capacity (Rux, Song et al. 2016). These studies typically rely on reducing the expression of one *Hox* paralogous group (or use single *Hox* knockout mouse models), however, and disregard the potential complementation by other highly expressed *Hox* genes in the cells compartment under study – as observed in the tibia periosteum (**Supplementary Fig. 1A, B**). The examination of compound mutants or deficiencies within a *Hox* cluster or paralogous group may therefore yield more severe phenotypes that can elucidate the role of *Hox* in adult skeletal cells. Here, our work shows that the expression of multiple posterior *Hox* paralogous groups must be decreased to detect a defect in number and function of tibial PSPCs; a reduction of part of the *Hox10* and *11* groups was not sufficient to reveal this phenotype (data not shown). This redundancy may represent an evolutionary mechanism to maintain stem cells.

Overall, the integration of these findings suggests a model in which, initially, the overlapping expression of several similar *Hox* paralogs collectively maintains skeletal stem cell function, after which individual *Hox* genes impart specific functions in the committed progenitor populations that direct their differentiation. This regulatory pattern would not be unprecedented in the context of *Hox*. In the developing Drosophila heart, forced expression of *Abd-B* (the orthologue of the posterior *Hox9-13* genes in mammals) inhibits cardiac myogenesis of mesodermal cells, underscoring its capacity to inhibit cell fate commitment in this tissue. Later during heart development, however, *Abd-B* expression is detectable in the more committed cells of the heart tube, precluding an inhibitory role at this later stage (Lovato, Nguyen et al. 2002).

In bone and other tissues, overlapping *Hox* genes show preferential activities that are consistent with a model of both simultaneous redundancy and specificity of *Hox* function (Shen, Montgomery et al. 1997, Shen, Rozenfeld et al. 1997, Pineault, Helgason et al. 2002, Akbas and Taylor 2004, Hedlund, Karsten et al. 2004). Thus, the observed co-expression of multiple *Hox* genes in periosteal cells highlight a probable complex combinatorial function in their regulation of primitive skeletal tissues. Further investigation of the transcriptional targets of HOX factors in distinct skeletal stem and progenitor cell populations may provide insight into their precise spatiotemporal and cell-type-specific roles.

The inability of *Hoxa10* overexpression to reprogram more committed PP2 progenitors is noteworthy. Cells that have progressed along the lineage trajectory and become fate-restricted likely undergo chromatin remodeling events that may inhibit HOX transcription factors from accessing the genes that regulate stem cell activity – enabling them to instead regulate differentiation genes as proposed above – but these hypotheses remain to be verified. Our previous work showed that the calvarial periosteum exhibits a near absence of *Hox* expression, contains more fate-restricted cells, and more inaccessible chromatin (Bradaschia-Correa, Leclerc et al. 2019). When we ectopically induced *Hox* expression in this cell population, we did not observe an upregulation of stem cell markers or an increase in calvarial PSPC number, as with tibial PSPCs (**Fig. 4, Supplementary Fig. 5**). This is in accordance with the notion that chromatin remodeling in more committed cells may prevent reprogramming by *Hox*.

The continued regional specification of *Hox* gene expression in adult tissues has been demonstrated by several independent studies, largely by the characterization of cells in culture (Chang, Chi et al. 2002, Leucht, Kim et al. 2008, Bradaschia-Correa, Leclerc et al. 2020). Here we corroborate this finding in the periosteal cell compartments of various anatomical regions and, importantly, find that this adult Hox code is functionally relevant to skeletal stem cell regulation. Region-specific *Hox* function in reprogramming and stem cell maintenance has implications for devising stem cell therapies that target specific segments of the skeleton, or potentially other tissues whose function is controlled by *Hox* expression profiles.

Relative to the bone marrow, the periosteum is a new, relatively understudied field of research. Periosteal progenitor cells play a central role in bone repair and, as such, represent a promising source of cells for tissue engineering approaches. PSPCs also exhibit a number of characteristics that are advantageous for such strategies, including their high proliferative rate necessary for efficient *in vitro* expansion (Sakaguchi, Sekiya et al. 2005, Yoshimura, Muneta et al. 2007, van Gastel, Torrekens et al. 2012), and a greater osteogenic capacity than many other mesenchymal stem cell populations both *in vitro* and when transplanted *in vivo* (Roberts, Geris et al. 2011, Roberts, van Gastel et al. 2015). Looking forward, advances in lineage reprogramming in many tissues have revealed a remarkable flexibility in cell identity (Morris 2016), and unraveling the mechanisms of this process in skeletal tissues can facilitate the development of cell fate engineering strategies. Further research examining how *Hox* overexpression increases stem cell potency along with the downstream genetic elements that mediate it – and investigating whether it does so without affecting lineage potential – can help achieve this therapeutic goal.

## Materials & Methods

### Animals

C57BL/6 mice, 8- to 16-week-old, were purchased from the Jackson Laboratory (Bar Harbor, ME) and bred in the barrier facility at the New York University School of Medicine. *Pdgfrα^CreERT/+^* knock-in mice were obtained from by the Michael Wosczyna Laboratory at NYU Langone (Wosczyna, Konishi et al. 2019). *Osx^CreERT2/+^* mice were received from Dr. H. M. Kronenberg, Massachusetts General Hospital. *Col1a1^CreERT/+^* mice (B6.Cg-Tg(Col1a1-cre/ERT2)1Crm/J) were obtained from JAX (016241). All mice were bred in the barrier facility at the New York University School of Medicine. To induce recombination in transgenic cre-ERT2 mice, tamoxifen (Sigma-Aldrich, St. Louis, MO, USA) was administered intraperitoneally either 2 mg/day according to the dosing protocol in **Figure 3A**. Mice were maintained on a 12-h light/dark cycle with food and water provided ad libitum.

### Patients and Specimens

All experiments involving human subjects were approved by the New York University (NYU) School of Medicine Institutional Review Board. After informed consent was obtained, bone marrow specimens were obtained during surgery at the fracture site. One cubic centimeter of bone marrow was immediately transferred into a microcentrifuge tube and placed on ice.

### Bulk RNA sequencing and Nanostring

FPKM values for each *Hox* gene was derived from tibial periosteal RNA sequencing data previously published by our group (Bradaschia-Correa, Leclerc et al. 2019) and CD45^−^ TER119^−^CD31^−^LEPR^+^ also published by our group (Josephson, Bradaschia-Correa et al. 2019). Nanostring^TM^ read counts were determined using the nCounter platform and by generating a custom panel of target-specific oligonucleotide probes (CodeSet) of the 39 murine *Hox* genes (**Table 3**). Of the total 78 Hox isoforms produced by the four Hox clusters, only one isoform of Hoxc4 was not detectable by the custom CodeSet. Five housekeeping genes (*Actb*, *Gusb*, *Pgk1*, *Tbp*, *Tubb*) were used to normalize the read counts.

**Table 3.**
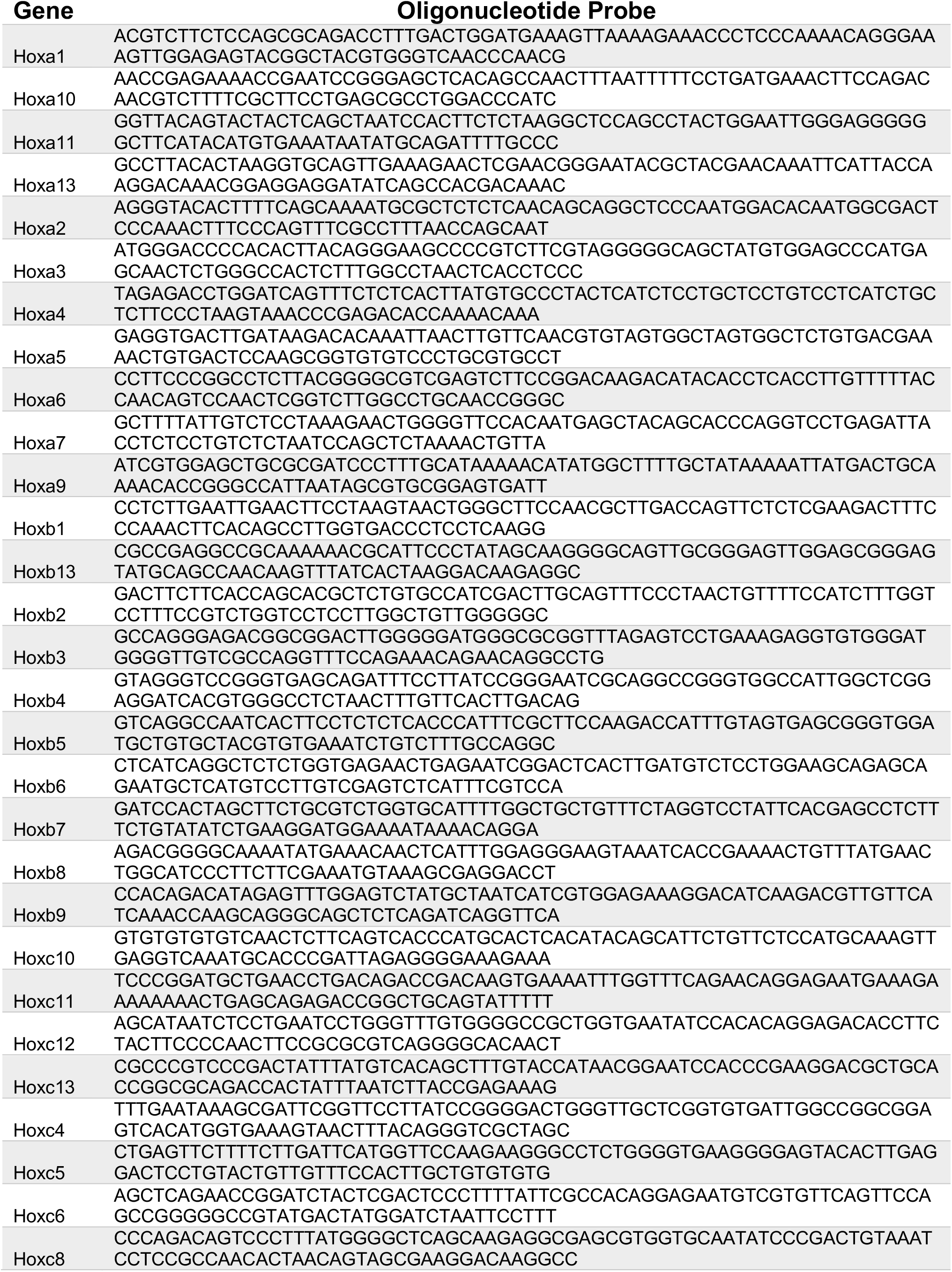
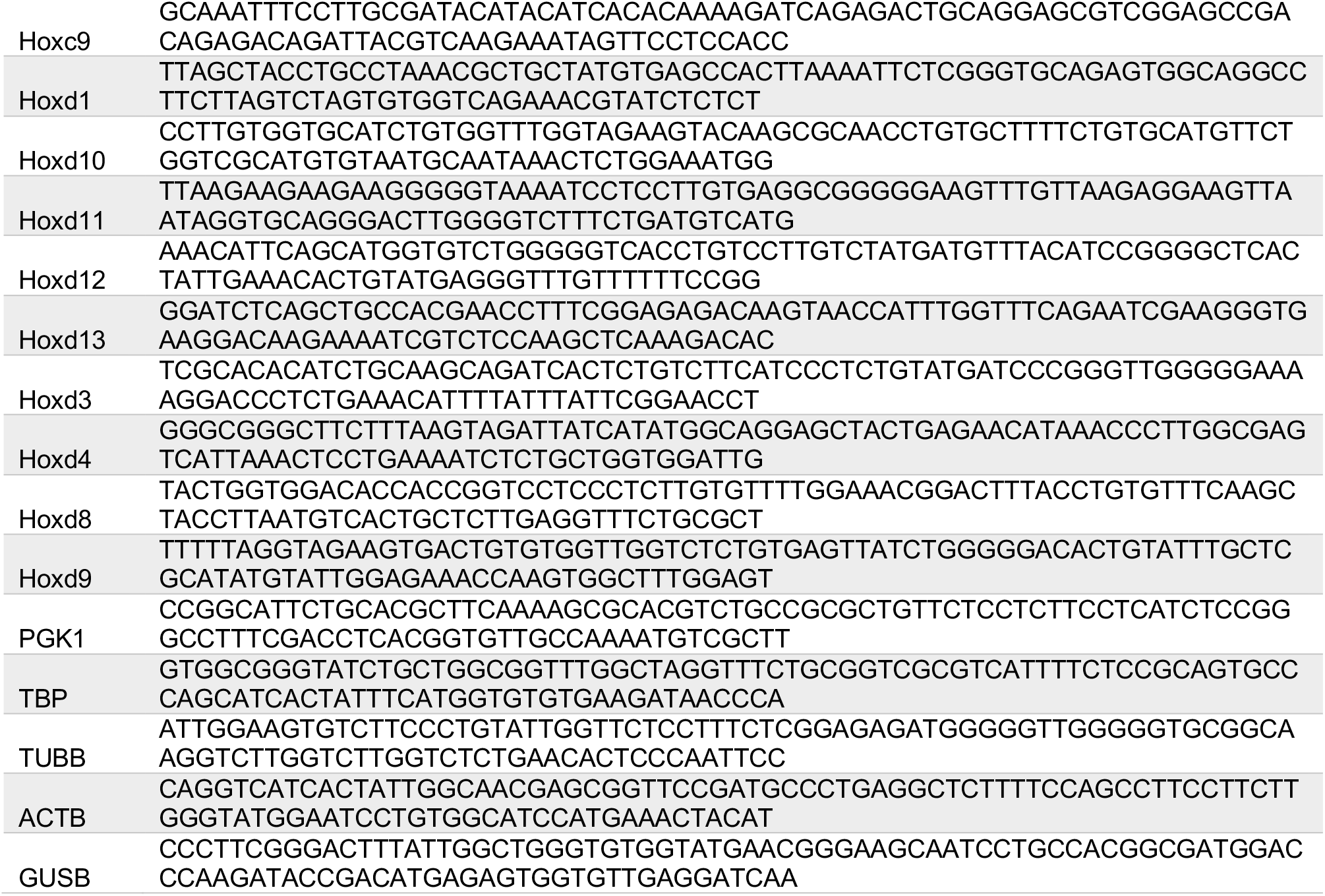
Nanostring^TM^ Custom Hox CodeSet. Oligonucleotides probes generated for the 39 Hox genes and 5 housekeeping genes used to probe absolute gene expression.

### Periosteal Cell Isolation

Primary periosteal stem and progenitor cells were obtained from the tibia, pelvis, anterior ribs (1-4), thoracic vertebrae (5-8) of the spine, radius/ulna, or parietal/frontal calvaria. After careful dissection from 8 to 16-week-old wild type (C57BL/6) mice, bones with intact periosteum were submitted to 4 serial collagenase digestions in 0.2% collagenase type 2 (ThermoFisher Scientific: 17101015) in DMEM (Life Technologies: 11885092) at 37 °C for 20 minutes with gentle rocking. After each of the first three digestions, bones were subjected to light centrifugation (1000 rpm) for 5 min and then transferred to a fresh tube of collagenase. After the last digestion, bones were centrifuged at 1400 rpm for 5 min and the pelleted cells were resuspended in growth media (GM): low glucose DMEM (Life Technologies: 11885092), 10% Fetal Bovine Serum (Life Technologies: 10437-028), 1% Penicillin/Streptomycin (Life Technologies: 15140122). Selective enrichment of periosteal stem/progenitor cells was confirmed using FACS analysis (**Supplementary Fig. 1C**).

### Flow Cytometry

Cells were trypsinized, resuspended in HBSS (Life Technologies: 14170161), supplemented with 2% Fetal Bovine Serum (Life Technologies: 10437-028), 1% Penicillin/Streptomycin (Life Technologies: 15140122), and 1% HEPES (Life Technologies: 15630080) (complete HBSS), and stained with 1:300 diluted CD45-PE (Miltenyi Biotec: 130-117-498), TER119-PE (Miltenyi Biotec: 130-117-512, TIE2-PE (ThermoFisher Scientific: 12-5987-82), 6C3-PE (ThermoFisher Scientific: 12-5891-82), CD90-PE (Invitrogen: MA5-17749), and 1:200 diluted CD51-BV421 (BD Biosciences: 740062), CD105-PE-Cy7 (ThermoFisher Scientific: 25-1051-82), CD200-BV711 (BD Biosciences: 745548) for 30 minutes on ice in the dark. Cells were then washed with 1mL of the complete HBSS solution, centrifuged at 1500 rpm for 5 minutes and finally resuspended with complete HBSS for flow cytometry. Cells were sorted on a Sony Biotechnology SY3200^TM^ cell sorter into a 50%/50% solution of complete HBSS and Fetal Bovine Serum or analyzed on a Bio-Rad ZE5 Analyzer. Sorting was validated to result in >95% purity of the intended population in postsort fractions. Beads (eBioscience 01-1111-41) were used to set initial compensation. Fluorescence minus one (FMO) controls were used for additional compensation and to assess background levels for each stain. We excluded doublets and gates were drawn as determined by internal FMO controls to separate positive and negative populations for each cell surface marker. Mesenchymal cell populations negative for CD45, CD31 and Ter119 cell surface markers were analyzed according to the approach described in Supplementary figure 2d.

### *in vitro* Differentiation

5 x 10^4^ PSPCs or sorted PP1 cells were seeded onto individual wells of 24-well plates (wells were first coated with a 1:100 dilution of fibronectin [Sigma: F0895] in PBS for 60 mins) in GM and allowed to attach overnight. The next day, cells were stimulated with osteogenic media (OM) [DMEM, 10% FBS, 100 μg/mL ascorbic acid, 10 mM β-glycerophosphate, and 1% penicillin/streptomycin]. Media was replenished every 2-3 days. For adipogenic differentiation, the cells were induced the next day using the MSC Adipogenic BulletKit (Lonza, Allendale, NJ) induction media.

### RNA Interference

Primary PSPCs were transfected with Qiagen’s commercially available GeneSolution siRNAs targeting *Hoxa10* (CAGGGCCCAGCCAAACTCCAA; SI00201859), *Hoxd10* (CCGAACAGATCTTGTCGAATA; SI00206542), *Hoxc10* (CAGGGCCCAGCCAAACTCCAA; SI00201859), *Hoxa11* (CACCACTGATCTGCACCCAAA; SI01068788), and *Hoxd11* (CCCGTCGGACTTCGCCAGCAA; SI01069558). AllStars Negative Control siRNA (Qiagen, 1027281) was used as a non-targeting control. Each component siRNA of *HoxMix* was delivered at 5μM, yielding a total *HoxMix* concentration of 25μM; non-targeting control siRNA was delivered at 25μM. HiPerfect Transfection Reagent (Qiagen) was used as a transfection reagent as per manufacturer’s instructions. Transfection was carried out at the moment of seeding onto multiwell plates before the cells fully attached to the plates (fast-forward transfection). The seeded cells were treated with siRNAs every 3 to 4 days and samples were assayed by qRT-PCR or flow cytometry after 2 to 14 days of knockdown.

### Proliferation Assay

5 x 10^4^ PSPCs were seeded onto wells of 24-well plates in GM and simultaneously administered either *HoxMix* or nontargeting siRNAs. After 24 hours, the cells were incubated with 10μM EdU at 37°C for 15 hours, washed with PBS, and then trypsinized. The Click-iT^TM^ Plus EdU Alexa Fluor^TM^ 488 Flow Cytometry Kit (ThermoFisher Scientific, C10632) was utilized to fix, permeabilize, and label EdU-incorporated cells with a Click-iT^TM^ reaction as per the manufacturer’s instructions before subjecting the cells to flow cytometry analysis thereafter.

### Cell-Cycle Analysis

CellTrace^TM^ Far Red Cell Proliferation Kit (ThermoFisher Scientific: C34564) was used per manufacturer’s instructions. In the RNA interference expreiments, PSPCs isolated from C57BL/6 wild-type mice were first expanded in vitro, and 5 x 10^4^ cells were then seeded onto individual wells of a 24-well plate with *HoxMix* or nontargeting siRNA. 24 hours later, the cells from each well were trypsinized, incubated with CellTrace^TM^ for 1 hour at 37°C on day 0, then replated and cultured for 6 days. On day 6, cells were trypsinized and stained for PDGFRα (Invitrogen: 25-1401-82). A separate batch of cells was also trypsinized and incubated with CellTrace^TM^ for 1 hour at 37°C on day 6 to serve as a positive control.

For the overexpression experiments, PSCs, PP1, and PP2 cells were sorted from *in vitro*-expanded PSPCs. 3 x 10^4^ cells were incubated with either *LV-GFP* or *LV-Hoxa10/GFP* in individual wells of a 24-well plate. After 24 hours, the procedure was then continued as described above. Cells in this case were stained with the *Debnath et al.*-defined lineage cell surface markers previously described.Cells were then analyzed on a BD Biosciences LSRII UV cell analyzer for dye dilution and surface marker profile.

### RNA Isolation and Quantitative Real-Time PCR

RNA was either isolated from cells immediately following periosteal isolation and FACS to observe *in vivo* gene expression or from cells expanded *in vitro*. The RNeasy Plus Kit (Qiagen: 74134) was used to isolate RNA and remove genomic DNA, and RNA was then reverse-transcribed with the iScript cDNA Synthesis Kit (Bio-Rad: 170-8891). Quantitative real-time PCR was carried out using the Applied Biosystems QuantStudio3 system and RT^2^ SYBR Green ROX PCR Master Mix (Qiagen: 330523). Specific primers were designed using Harvard PrimerBank (http://pga.mgh.harvard.edu/primerbank/) (**Table 4**). Results are presented as 2^−ΔΔCt^ values normalized to the expression of *18s*. Means and SEMs were calculated in GraphPad Prism 7 software.

**Table 4.**
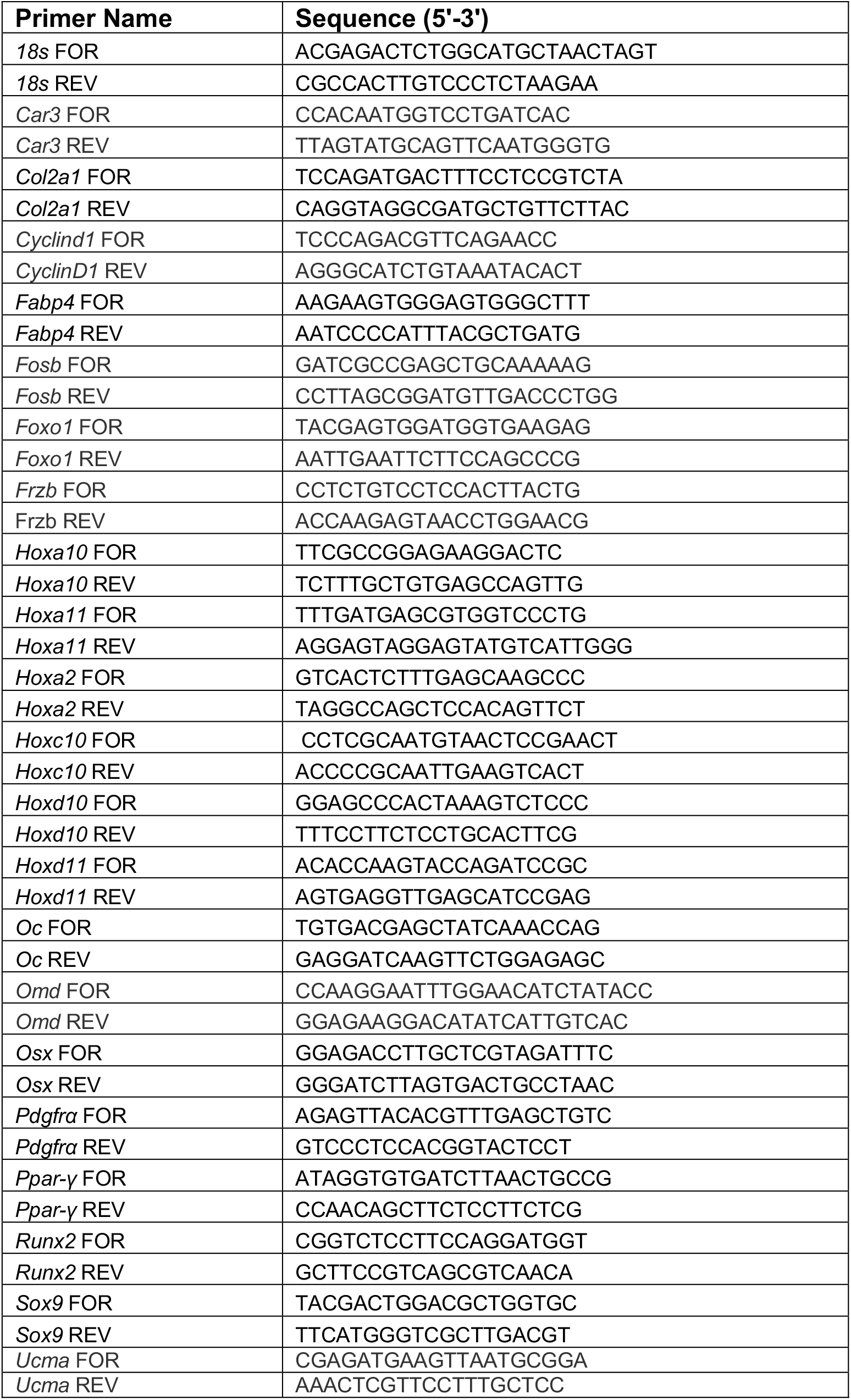
qRT-PCR primers. All primers were purchased from Integrated DNA Technologies.

### Viral Generation and Transduction

Lentiviral DNA containing either a *Hoxa10* CDS expression construct or a control construct lacking *Hoxa10* CDS was generated at Genewiz (New Jersey) using the Tet-ON system. In addition to the Hoxa10 sequence, the lentiviral vector used (*ptetO*) also include EGFP, Luciferase, and Puromycin cloned downstream of the active CMV promoter. 2A peptide sequences are also included between each element (*ptetO-Hoxa10-T2A-EGFP-P2A-Luciferase-T2A-Puromycin*; *LV-Hoxa10/GFP*) (*ptetO-EGFP-P2A-Luciferase-T2A-Puromycin*; *LV-GFP*) in order to produce multi-cistronic, equimolar expression of all four genes. EGFP was used to track the cells that have been infected in culture. Identical methods were used to generate lentiviral sequences containing *Hoxa11*, *Hoxb8*, and *Hoxa5*. *pLenti-rtTA3* (Addgene: 26429), *pRSV-Rev* (Addgene: 12259), *pMD2.G* (Addgene: 12259), and *pMDLg/pRRE* (Addgene: 12251) plasmids were purchased and lentivirus was generated in the Lenti-X™ 293T Cell Line (TakaraBio: 632180), purified with a Lenti-X™ Maxi Purification Kit (TakaraBio: 631234), and titered with a Lenti-X qRT-PCR Titration Kit (TakaraBio: 631235).

3 x 10^4^ sorted PSCs, PP1, PP2, or *in vitro*-expanded PSPCs were seeded onto individual wells of 24-well plates (wells were first coated with a 1:100 dilution of fibronectin [Sigma: F0895] in PBS for 60 mins) using GM made with 10% heat-inactivated Fetal Bovine Serum and 1% Penicillin/Streptomycin. The cells were immediately transduced with *pLenti-rtTA* and either *LV-GFP*, *LV-Hoxa10/GFP*, *LV-Hoxa11/GFP*, *LV-Hoxb8/GFP*, or *LV-Hoxa5/GFP* at an M.O.I. of 75. The transduction efficiency was aided by the addition of 2.5 μg/mL Polybrene (Sigma) and the cells were treated 10 μg/mL Doxycycline to activate expression downstream of the tetO promoter sequences. GM with 10 μg/mL Doxycycline was used to replace the media every 2-3 days.

### Identification of PSC Markers from Dataset

The periosteal scRNA-seq dataset is from a publicly available adult mouse femoral periosteum study (Debnath, Yallowitz et al. 2018). We obtained the raw count matrix from the GEO accession ID <u>GSE106236</u> and annotated cells based on high expression levels of the genes associated with the cell surface markers used for flow cytometry [6C3 (*Enpep*), CD90 (*Thy1*), CD51 (*Itgav*), CD105 (*Eng*), and CD200 (*Cd200*)]. The PSC, PP1, and PP2 cells in the count matrix were then sorted *in silico* according to the *Debnath et al.*-derived gating strategy presented in the results, and genes with a high fold change between PSCs and PP1/PP2 cells were used to identify potential PSC markers.

### Renal Capsule Transplants

A model of mesenchymal stem cell differentiation was used to compare the regenerative potential of PSPCs. 12- to 15-week-old, syngeneic C57BL/6 mice were used as hosts for the renal capsule transplantation assay. An incision was made on the dorsal skin surface, followed by an incision through the peritoneum, the kidneys were identified and then exteriorized. An incision in the renal capsule was made using a 27-gauge needle. Two microliters of tibial bone marrow (containing ∼100,000 cells) from 12-week-old C57BL/6 mice were used to resuspend 750 transduced (GFP^+^) PP1 cells. The mixture was then left exposed to open air for one to two minutes to allow for a limited amount of coagulation and subsequently grafted beneath the capsule. The kidney was placed back into its anatomic location and the peritoneum was closed with a Vicryl suture, followed by skin closure with a 6-0 Vicryl suture. Mice had ad lib access to food and water (with dissolved .4 mg/mL Doxycycline and 5% sucrose) and received subcutaneous buprenorphine for analgesia. Mice were euthanized 14 days post-surgery, the renal grafts were harvested, digested in 0.2% collagenase type 2 (ThermoFisher Scientific) in DMEM at 37°C for 1 hour, stained with antibodies, and subjected to FACS. The GFP^+^ cells were collected and reused for the subsequent renal grafts.

---

**Supplementary Figure 1.**
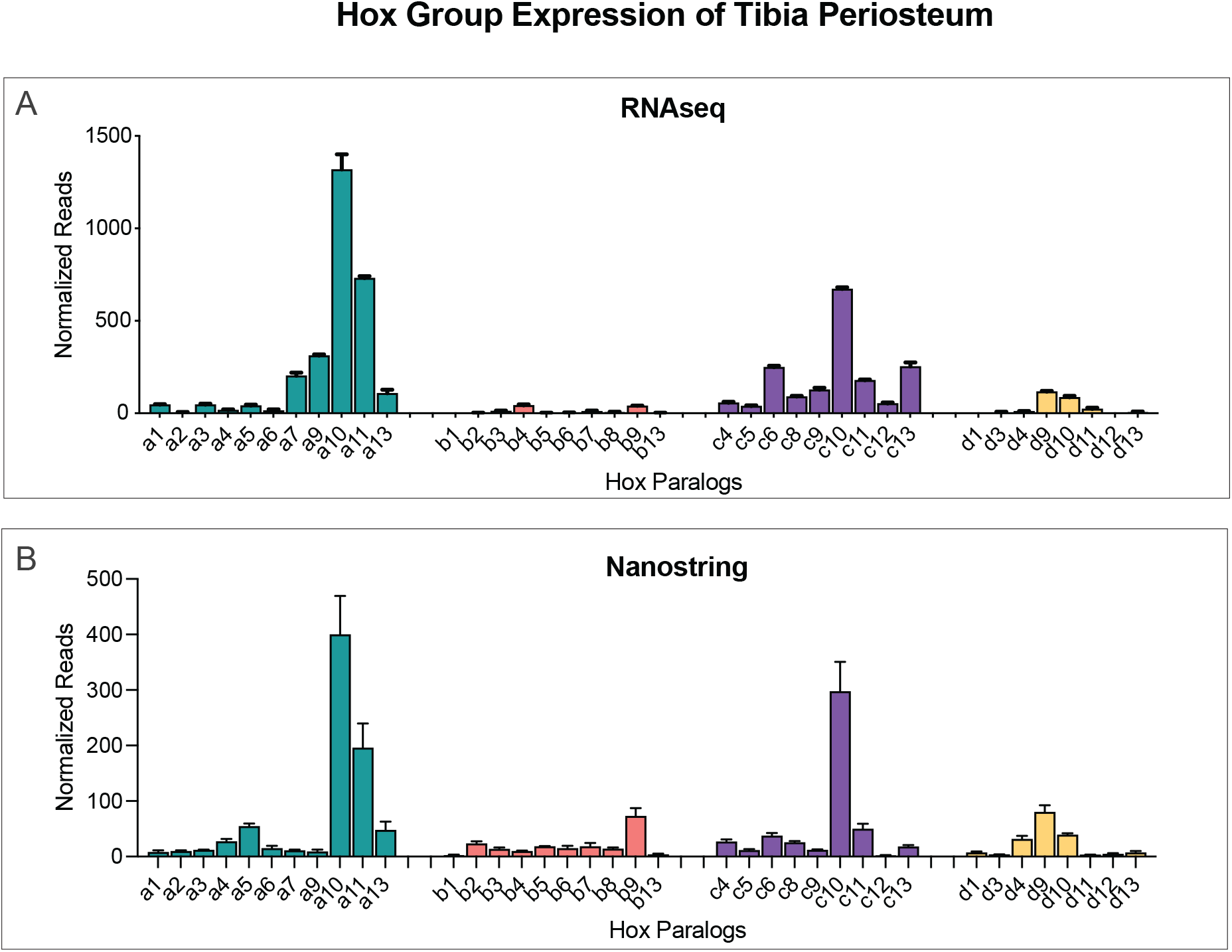
*Hox* expression profile of adult tibia periosteum. (**A**). Bulk RNAseq revealed the gene expression pattern for all 39 Hox genes in 12-week-old, freshly-isolated tibial periosteum. *Hoxa10*, with the most normalized reads, is the most highly expressed. *n* = 3 mice. (**B**) Nanostring^TM^ probes against all 39 Hox genes revealed their absolute expression profile in the adult tibia periosteum. *Hoxa10* contained the most normalized reads. *n* = 4 mice. (**C**) Representative FACS plot demonstrating the enrichment in PSPCs following 10 days of *in vitro* expansion of cells isolated from the periosteum. Error bars are SEM.

**Supplementary Figure 2.**
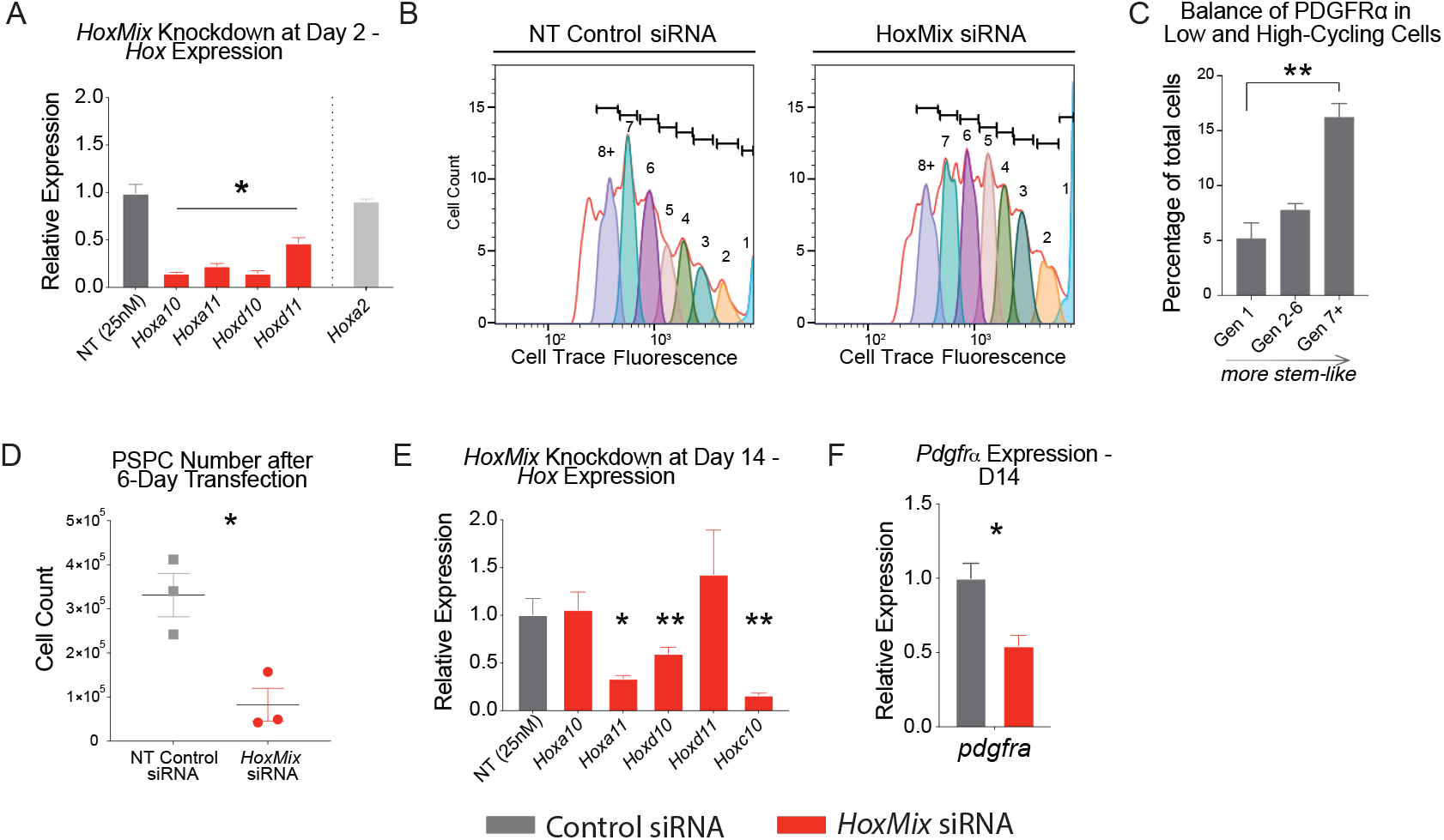
Knockdown of posterior *Hox* genes in adult tibial PSPCs. (**A**) The expression of several posterior *Hox* genes in cultured PSPCs two days after knockdown with 5nM each of si*Hoxa10*, si*Hoxa11*, si*Hoxd10*, si*Hoxd11*, and si*Hoxc10* (termed *HoxMix* siRNA) normalized to their corresponding expression when PSPCs are transfected with non-targeting (NT) control siRNA (25nM). The expression of *Hoxa2* was used to ascertain specificity of the posterior *Hox* siRNAs. *n* = 3. (**B**) Representative cytometric plots of siControl- and si*HoxMix*-transfected tibial PSPCs that were treated with Cell Trace^TM^ to categorize cells by cell cycle generation (from 1 to 8+) after six days of incubation. *n* =5 (control), *n* =7 (*HoxMix*). (**C**) The distribution of PSPCs displaying the skeletal stem cell marker, PDGFR*α*, among low-cycling (generation 1), medium-cycling (generation 2-6), and high-cycling (generation 7+) cells as measured by flow cytometry. (**D**) The absolute number of PSPCs after an equal seeding density and 6 days of transfection with either NT control or *HoxMix* siRNA. *n* = 3. (**E**) The expression of several posterior *Hox* genes in cultured PSPCs 14 days after a serial knockdown with 25nM of si*HoxMix* normalized to their corresponding expression in PSPCs transfected with 25nM NT control siRNA. *n* = 3. (**F**) qRT-PCR measured the relative gene expression of *pdgfrα* 14 days after a serial knockdown of PSPCs with si*HoxMix and* siControl. *n* = 3. \**p* < 0.05, \*\**p* < 0.01. Two tailed Student’s t-test. Error bars are SEM.

**Supplementary Figure 3.**
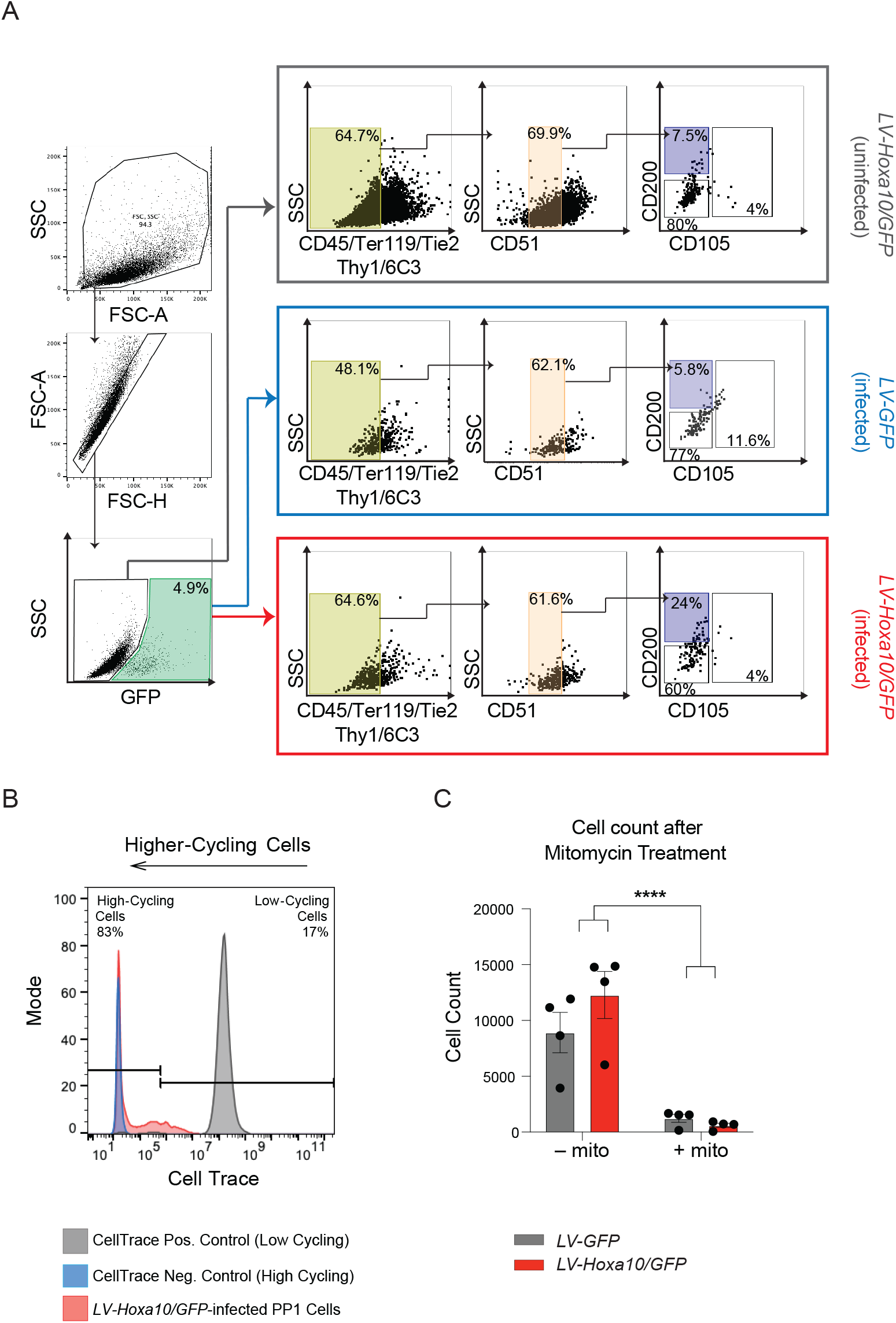
Hoxa10 overexpression mediates reprogramming of adult tibial PP1 cells into PSCs. (**A**) Representative FACS plots of the PSPC lineage output 7 days after transduction of PP1 cells with *LV-GFP* or *LV-Hoxa10/GFP*. Infected (GFP^+^) and uninfected (GFP^−^) cells were evaluated separately. (**B**) Flow cytometric analysis on a CellTrace^TM^ positive control (dye administration 30 minutes before analysis) and on a CellTrace^TM^ negative control (no dye administration) was used to categorize cells as low- or high-cycling using the indicated gating strategy. A sample of *LV-Hoxa10/GFP*-infected PP1 cells after six days of *in vitro* administration of CellTrace^TM^ was used as a representative sample. (**C**) The number of cells seven days after transduction of PP1 cells with LV-GFP or LV-Hoxa10/GFP with or without 10 μg/μl Mitomycin C treatment2. *n* = 4. \*\*\*\**p* < 0.0001. Two tailed Student’s t-test. Error bars are SEM.

**Supplementary Figure 4.**
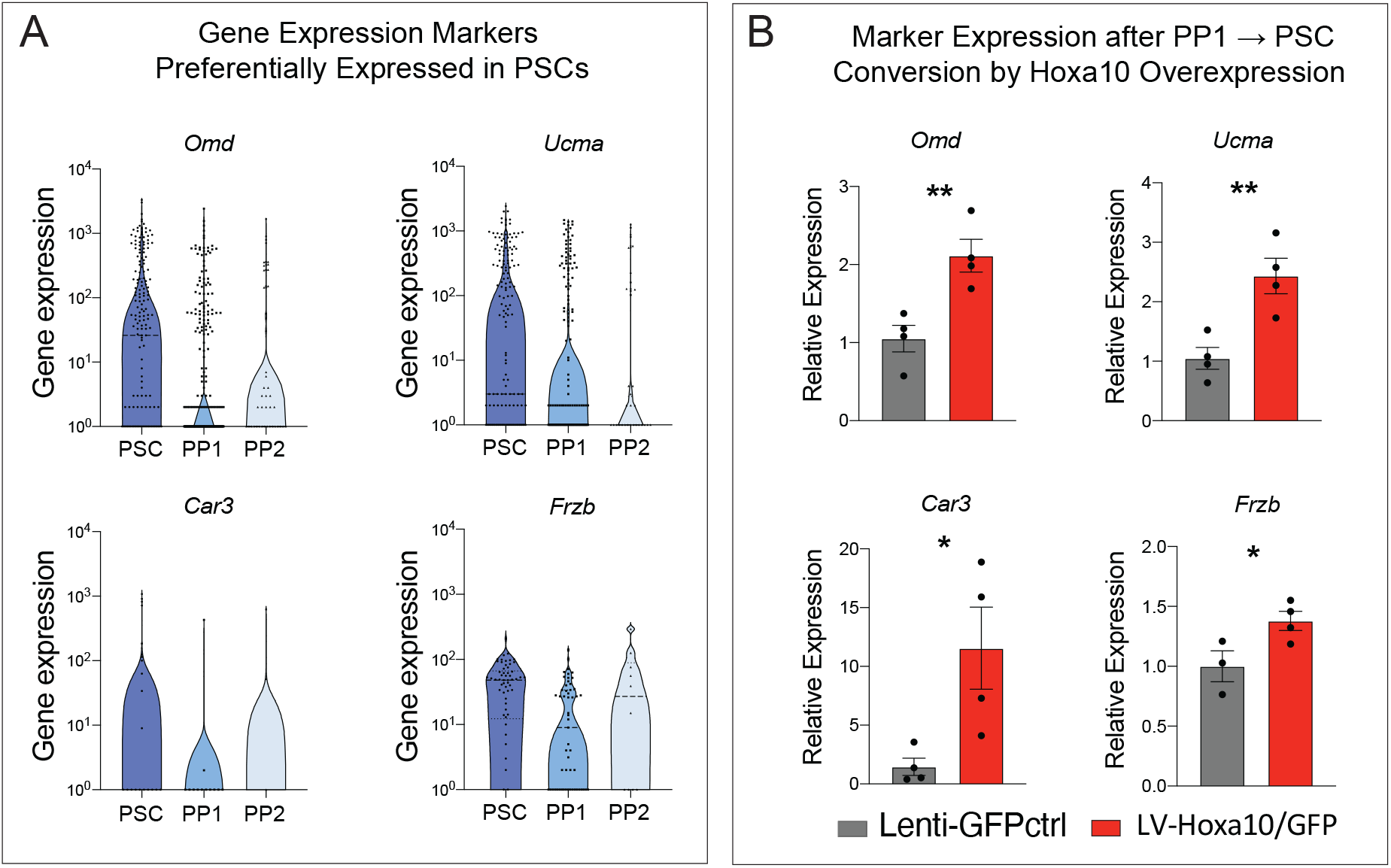
Enriched periosteal stem cell marker expression after reprogramming of tibial progenitors. (**A**) Single-cell RNAseq data from Debnath, Yallowitz et al. 2018 was analyzed to obtain a list of marker genes that are enriched in PSCs relative to PP1 cells. PSC, PP1, and PP2 cells were sorted *in silico* based on genetic expression of CD51 (*itgav*), CD105 (*eng*), and CD200 (*cd200*) and on the gating strategy presented in Debnath, Yallowitz et al. 2018. *n* = 205 PSCs, *n* = 393 PP1 cells, *n* = 60 PP2 cells. (**B**) Relative gene expression of PP1 cells seven days after transduction with *LV-GFP* or *LV-Hoxa10/GFP* for the previously defined PSC markers. *n* = 4, *LV-GFP*; *n* = 4, *LV-Hoxa10/GFP.* \**p* < 0.05, \*\**p* < 0.01. Two tailed Student’s t-test. Error bars are SEM.

**Supplementary Figure 5.**
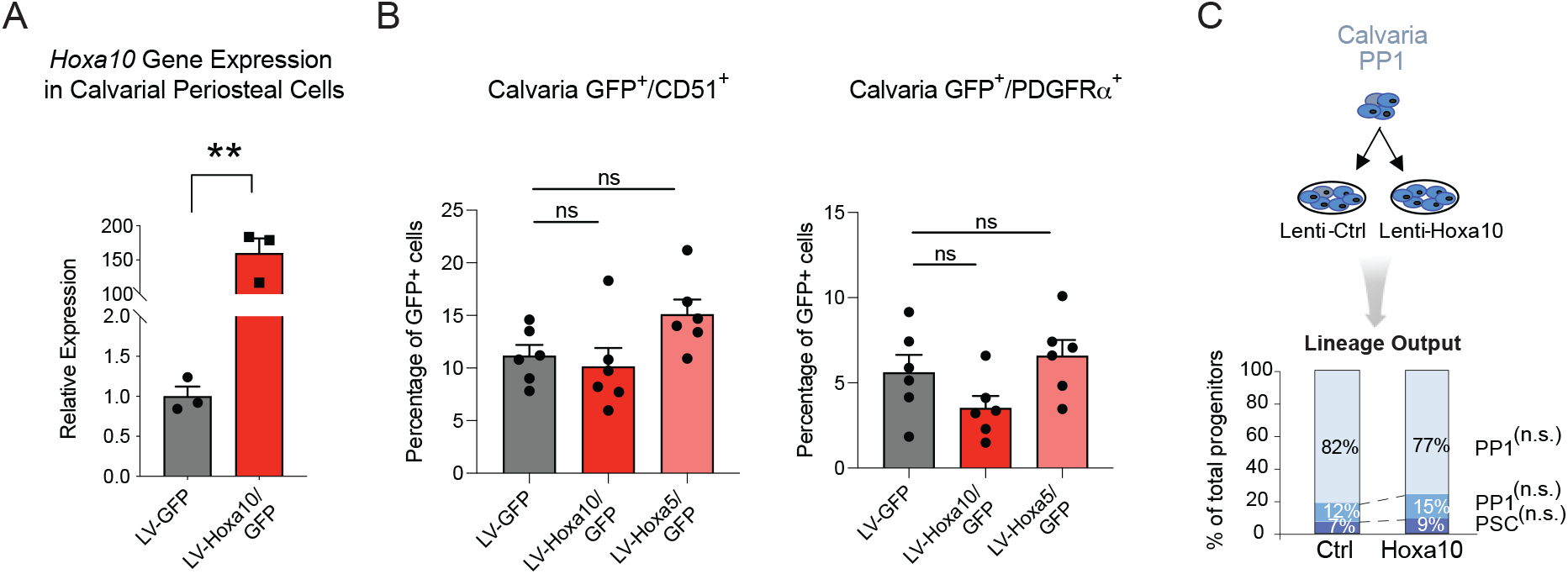
*Hoxa10* overexpression in calvarial periosteal cells does not increase stem cell marker expression. (**A**) Relative expression of *Hoxa10* and *Pdgfrα* in calvarial PSPCs 6 days after infection with *LV-GFP* or *LV-Hoxa10/GFP,* as measured by qRT-PCR*. n* = 3, *LV-GFP*; *n* = 3, *LV-Hoxa10/GFP* (**B**) Seven days after transduction, flow cytometry revealed the balance of CD51^+^, PDGFR*α*^+^, or SCA1^+^ stem cells among *LV-GFP*- or *LV-Hoxa10/GFP*-infected calvarial PSPCs. *n* = 3, *LV-GFP*; *n* = 2, *LV-Hoxa10/GFP.* \*\**p* < 0.01. Two tailed Student’s t-test. Error bars are SEM.

**Supplementary Figure 6.**
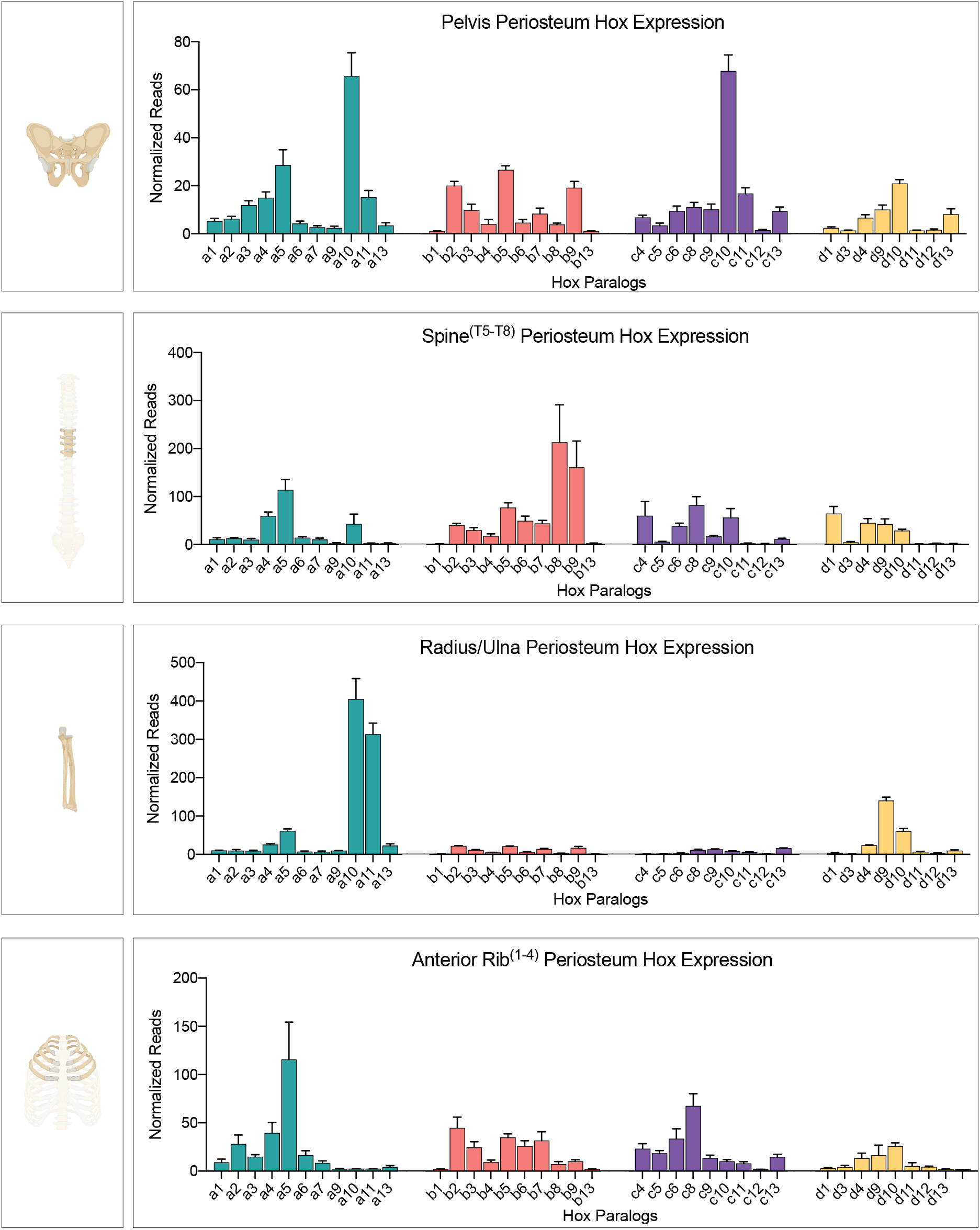
*Hox* expression profile of 12-week-old periosteum from various anatomical regions. Nanostring^TM^ probes against all 39 Hox genes revealed their absolute expression profile in freshly-isolated periosteum of the pelvis, spine^T5-T8^, radius/ulna, and anterior rib^1-4^. *n* = 4 mice. Error bars are SEM.

**Supplementary Figure 7.**
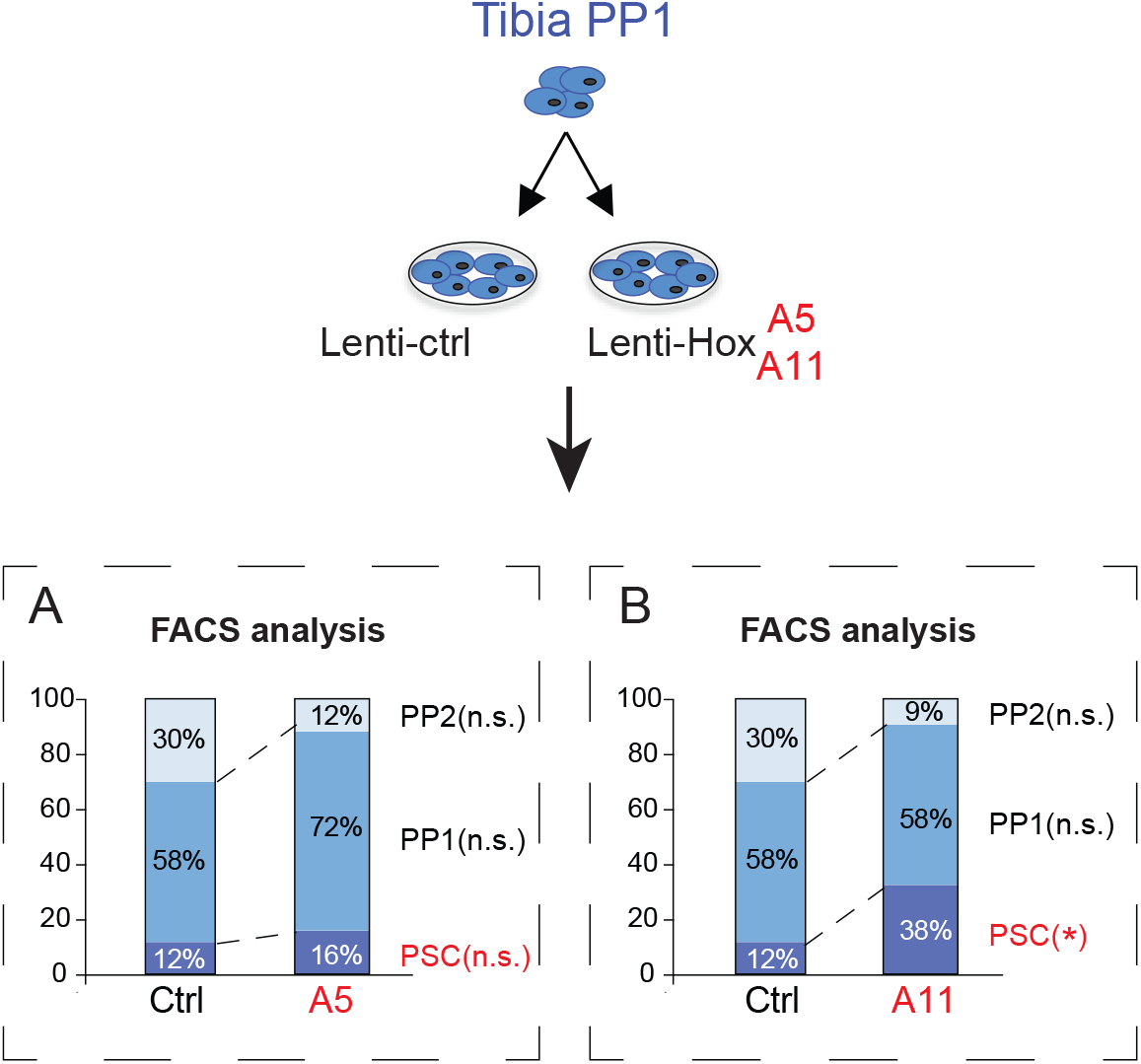
Reprogramming of tibial periosteal progenitors is *Hox* code-dependent. (**A**) *Hoxa5*, an anterior *Hox* gene, does not change the lineage output of tibial progenitor cells 7 days after transduction with *LV-GFP* (Ctrl) or *LV-Hoxa5/GFP* (A5). (**B**) *Hoxa11* (adjacent to *Hoxa10*) shifts the lineage output of tibial progenitor cells 7 days after transduction with *LV-GFP* (Ctrl) or *LV-Hoxa11/GFP* (A11). Full lineage output data and statistics are provided in Table 2. *n.s.* = not significant, \**p* < 0.05. Two tailed Student’s t-test.

